# Cell type-specific disruption of cortico-striatal circuitry drives repetitive patterns of behaviour in fragile X syndrome model mice

**DOI:** 10.1101/2022.12.08.519509

**Authors:** Francesco Longo, Sameer Aryal, Paul Anastasiades, Marta Maltese, Corey Baimel, Federica Albanese, Joanna Tabor, Jeffrey D Zhu, Mauricio M Oliveira, Denise Gastaldo, Claudia Bagni, Emanuela Santini, Nicolas X Tritsch, Adam G Carter, Eric Klann

## Abstract

Individuals with fragile X syndrome (FXS) are frequently diagnosed with autism spectrum disorder (ASD), including increased risk for restricted and repetitive behaviours (RRBs). Consistent with observations in humans, FXS model mice display distinct RRBs and hyperactivity that are consistent with dysfunctional cortico-striatal circuits, an area relatively unexplored in FXS. Using a multidisciplinary approach, we dissected the contribution of two populations of striatal medium spiny neurons (SPNs) in the expression of RRBs in FXS model mice. We found that dysregulated protein synthesis at cortico-striatal synapses is a molecular culprit of the synaptic and ASD-associated motor phenotypes displayed by FXS model mice. Cell-type-specific translational profiling of the FXS mouse striatum revealed differentially translated mRNAs, providing critical information concerning potential therapeutic targets. Our findings represent the first evidence of a cell-type specific impact of the loss of FMRP on translation and the sequence of neuronal events in the striatum that drive RRBs in FXS.

**Highlights:** Dysregulated striatal protein synthesis underlies altered synaptic plasticity and RRBs displayed by FXS model mice

FXS model mice exhibit cell type-specific molecular, morphological, and synaptic changes in the dorsolateral striatum

Selective deletion of *Fmr1* from dSNPs in mice alters translation and causes repetitive behavior

TRAP-Seq indicates that there is altered binding of >120 mRNAs to ribosomes in dSNPs of *Fmr1* KO mice

G-protein signaling (RGS) 4 translation is significantly reduced in dSPNs in FXS model mice

Treatment of FXS model mice with VU0152100, a positive allosteric modulator of the M4 muscarinic receptor that is upstream of RGS4, can reverse RRBs

## Introduction

Restricted and repetitive patterns of behavior and interest (RRBs) are one of the core symptoms that define autism spectrum disorder (ASD). They comprise a wide range of motor, cognitive, and behavioral traits that are manifested in a variety of combinations and levels of severity in individuals with ASD. RRBs can arise during infancy and are the first signs of ASD to emerge in toddlers^1,2^, whereas the persistence of RRBs during adolescence and adulthood often results in a barrier to learning and social interactions in ordinary life. However, despite the significant impact on both familial and social dynamics, the neural underpinnings of RRBs in ASD remain poorly understood.

Evidence from both clinical and preclinical studies strongly suggest that the expression of RRBs, as well as other ASD-associated behaviors such as cognitive inflexibility and impulsive/compulsive behavior, arise from altered cortico-striatal-thalamic-cortical circuitry^3,4^. As the main input nucleus to the basal ganglia, the striatum is directly engaged in the control of goal-directed actions and habits^5^. Striatal function relies on two distinct populations of GABAergic striatal spiny projection neurons (SPNs): direct pathway SPNs (dSPNs; expressing the dopamine D1 receptor) and indirect pathway SPNs (iSPNs; expressing the dopamine D2 receptor) that either promote or suppress action selection, respectively^5^. The heterogeneity of RRBs in ASD may mirror specific perturbations among the complexity of striatal circuitry. Both anatomical and functional studies indicate consistent structural alteration in striatal volume in ASD individuals^6^, which are associated with atypical striatal development^7^ as well as aberrant patterns of connectivity between the striatum and different ASD-relevant cortical areas^8,9^.

Individuals with fragile X syndrome (FXS), the most common form of inherited intellectual disability (ID), exhibit a variety of behaviors emblematic of ASD, including stereotypy, impaired social interaction, and anxiety^10^. FXS is associated with increased risk for RRBs including hand flapping, body rocking, self-injury, and compulsive behavior^2,11,12^. Neuroimaging and surface-based modelling studies have shown structural changes in the corpus callosum and putamen of FXS individuals, where the enlarged caudate nucleus positively correlates with reduced intellectual abilities and increasing levels of RRBs^13–16^. Despite the fundamental contribution of cortico-striatal circuit dysfunction in the development of RRBs, hyperactivity, and impaired social interaction in FXS, the effect of transcriptional silencing of the *FMR1* gene and the loss-of-function of its product fragile X messenger ribonucleoprotein (FMRP) in the striatum remains largely unexplored.

Dysregulated protein synthesis has emerged as a shared molecular anomaly that underlies the structural and functional synaptic plasticity impairments and aberrant behaviors associated with both FXS and ASD^17,18^. The impact of the loss of FMRP, an mRNA-binding protein that in most cases operates as a negative regulator of translation^19^, among other functions^19,20^, has been extensively explored in the hippocampus and cortex^19,21–24^. Evidence from both cells derived from FXS patients and preclinical models of FXS suggests that FMRP functions by blocking both initiation and elongation steps of translation and as a result of its absence, overall protein synthesis is enhanced^19,21^. During initiation, FMRP interacts with cytoplasmic FMRP-interacting protein 1 (CYFIP1), which associates and sequesters the cap-binding protein eukaryotic initiation factor 4E (eIF4E), thereby blocking its interaction with the eukaryotic initiation factor 4G (eIF4G) and inhibiting the translation of specific transcripts^25^. In addition to its direct action in repressing translation, FMRP regulates protein synthesis indirectly by suppressing the translation of components of the mammalian target of rapamycin complex 1 (mTORC1) signaling pathway^26^. In the absence of FMRP, the homeostatic balance that translational repression would have on the appropriate rate of local protein synthesis in response to synaptic activity is perturbed. As a result, many forms of long-term synaptic and spine morphological plasticity are altered in the cortex and the hippocampus of FXS model mice^19^, but the impact on striatal circuits remains to be examined.

Here, we adopted a multidisciplinary approach to investigate the molecular and synaptic mechanisms of cortico-striatal circuit dysfunction underlying the expression of RRBs and hyperactivity in mouse models of FXS. Our findings add to the emerging literature on RRBs in ASD by contributing the first evidence of cell-type specific changes in translation resulting from the loss of FMRP and the sequelae of neuronal events within the striatum that drive RRBs in FXS.

## Results

### *Fmr1* KO mice display facilitation of locomotor activity and engage in repetitive/perseverative behaviors

To investigate the role of FMRP in overall motor ability and RRBs resembling those observed in humans with FXS^2,11,12^, we tested *Fmr1* knockout (KO) mice and their wild-type (WT) littermates on several different motor skill assays. Mice were tested for spontaneous horizontal and vertical activity in the open field (OF) and cylinder tests, respectively, as well as for novelty-induced activity in the novel home cage (NHC) tests. In addition, the drag and pole tests were used to assess both bradykinesia and motor coordination. *Fmr1* KO mice displayed an overall hyperactive motor phenotype (**Fig. 1a-e**). In the OF test, *Fmr1* KO mice exhibited significantly greater distance travelled than WT mice (**Fig 1a**). *Fmr1* KO mice also exhibited an increase in novelty-induced locomotor activity compared with their littermate controls (**Fig. 1b**). Consistent with the motor facilitation exhibited in both the OF and the NHC tests, *Fmr1* KO mice displayed significantly enhanced vertical locomotor activity in the cylinder test compared with their controls (**Fig. 1c**). Moreover, *Fmr1* KO mice exhibited significantly greater motor ability than WT controls in both the drag (**Fig. 1d**) and the pole test (**Fig. 1e**), motor tests specific for assessing striatal-driven locomotor activity.

**Figure 1:**
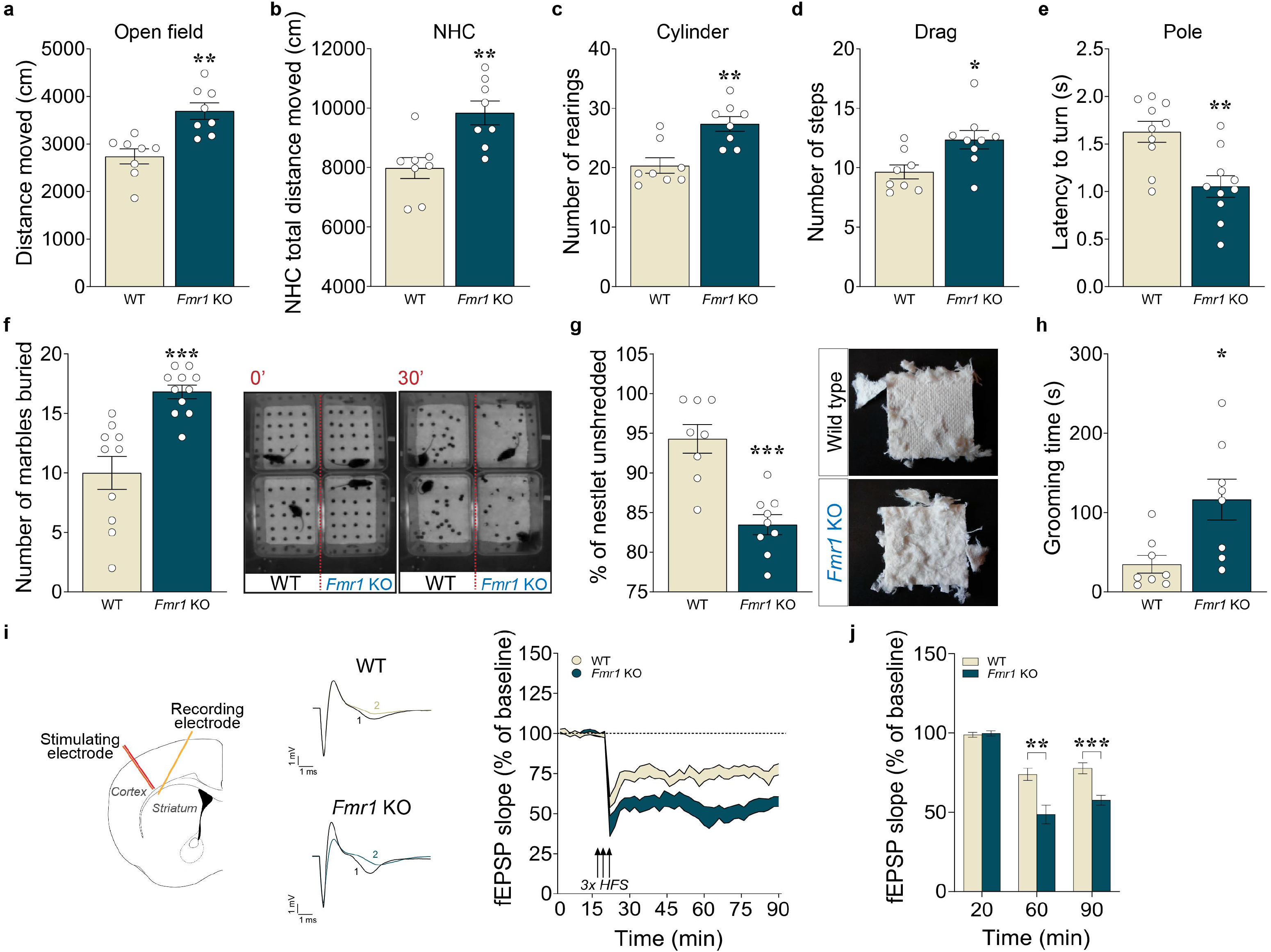
*Fmr1* KO mice exhibit increased locomotor activity, repetitive and perseverative behaviour, and altered cortico-striatal synaptic plasticity. *Fmr1* KO mice and their WT littermates mice were subjected to a set of tests for assessing global locomotor activity (**a-e**), including the open field (**a**), novel home cage (**b**), cylinder (**c**), drag (**d**) and, pole (**e**) tests. *Fmr1* KO mice showed increased locomotor activity compared to WT mice (**a-c**). **a**, Summary plot of spontaneous locomotor activity expressed as distance moved (cm) during the open field test over 15 min. (unpaired *t* test, t*_(14)_*= 4.06, \*\**P* <0.01; *n*=8 mice/genotype). **b**, Summary plot of the novelty-induced locomotor activity expressed as a novel home cage (NHC) distance moved (cm) in a 60 minutes test during novel home cage test (unpaired *t* test, t*_(14)_*= 3.51, \*\**P* <0.01; *n*=8 mice/genotype). **c**, Summary plot of vertical activity expressed as number of rearing episodes (number of counts) during the cylinder test over 5 min (unpaired *t* test, t*_(14)_*= 3.91, \*\**P* <0.01; *n*= 8 mice/genotype). *Fmr1* KO showed motor facilitation in motor tests specific for assessing striatal-based locomotor activity (**d-e**) when compared to their littermates control mice. **d**, Summary plot of average number of steps during drag test (unpaired *t* test, t*_(15)_*= 2.70, \**P* <0.05; *n*= 8-9 mice/genotype). **e**, Summary plot of latency to turn (s) during pole test (unpaired *t* test, t*_(18)_*= 3.60, \*\**P* <0.01; *n*= 10 mice/genotype). *Fmr1* KO exhibited exaggerated repetitive and perseverative behaviors compared to their WT control mice in the marble burying (**f**), nestlet shredding (**g**) and grooming (**h**) tests. **f**, Summary plot of number of marbles buried and representative image from video recorded before and after the 30 min marble burying test (unpaired *t* test, t*_(19)_*= 4.68, \*\*\**P* <0.001; *n*= 10-11 mice/genotype). **g**, Summary plot of percentage of unshredded nestlet during the nestlet shredding test (unpaired *t* test, t*_(15)_*= 4.99, \*\*\**P* <0.001; *n*= 8-9 mice/genotype). Mice were analyzed using Student’s *t* test. \**P* < 0.05, \*\**P* < 0.01, \*\*\**P* < 0.001 *Fmr1* KO mice different from age-matched WT littermates. **h**, Summary plot of time spent grooming in the last 10 minutes interval of a 60 minutes test (unpaired *t* test, t*_(14)_*= 2.90, \**P* <0.05; *n*= 8 mice/genotype). All data are shown as mean ± s.e.m. *Fmr1* KO mice exhibit enhanced cortico-striatal long-term depression (LTD; **i,j**). **i** (left panel), Schematic representation of electrode placement and representative traces of superimposed fEPSPs (scale bars represent 1 mV/ms) recorded during baseline (1) and 60 min after high frequency stimulation (HFS) train (2). Arrows indicate delivery of HFS. **i** (right panel), Plot showing normalized fEPSP mean slope (±s.e.m. displayed every 2□min) recorded from cortico-striatal slices. **j**, Mean fEPSPs at baseline (20 min), at 60 (40 min after tetanus) and at 90 min (70 min after tetanus). LTD evoked by 3 trains of HFS was significantly enhanced in *Fmr1* KO cortico-striatal slices at both 60 min and 90 min (two-way RM ANOVA, followed by Bonferroni’s multiple comparisons test, time x genotype, F*_(2,54)_* = 8.51, P = 0.0006; *n*=12-15 slices from 8 mice/genotype). Data are shown as mean ± s.e.m. \*\**P* < 0.01 and \*\*\**P* < 0.001 Fmr1 KO *versus* WT mice.

We next evaluated cohorts of *Fmr1* KO mice and their WT littermates for expression of core features of RRBs, which are a defining trait of ASD^1,2,27^. *Fmr1* KO and WT mice were tested in the marble-burying (MB) and nestlet-shredding tests as complementary methods for assessing repetitive behaviors in mice. *Fmr1* KO mice buried a greater number of marbles compared to controls (**Fig. 1f**). Likewise, *Fmr1* KO mice shredded significantly more of their nestlets compared to WT mice (**Fig. 1g**). The dorsolateral region of the striatum also is critical for the execution of normal grooming behavior^28^; thus, *Fmr1* KO mice and their littermate controls also were tested for self-grooming behavior. Consistent with the repetitive behavior revealed in both the MB and the nestled shredding tests, *Fmr1* KO mice engaged in significantly more grooming activity compared to their WT controls (**Fig. 1h**). Taken together, these results support and complement previously reported preclinical and clinical studies on RRBs^9,27^, and demonstrate that the lack of FMRP results in increased motor activity and the development of stereotyped/RRBs in mice.

### *Fmr1* KO mice exhibit a net increase in cap-dependent translation via increased eIF4E-eIF4G interactions, which contributes to altered synaptic plasticity, function, and spine density in DLS

Altered *de novo* protein synthesis is a shared phenotype between several preclinical models of ASD, including FXS mice^17–19,21,29^. Modifications of *de novo* translation stemming from the lack of FMRP expression have been widely investigated in different rodent brain areas^21,30–32^. However, few studies have focused on brain structures outside of the cortex and hippocampus, leaving the striatum largely unexplored. Given the significant changes in locomotor activity and the expression of RRBs by *Fmr1* KO mice (**Fig. 1**), all of which are influenced by striatal activity^4,27^, we first sought to examine whether the loss of *Fmr1* results in specific synaptic aberrations in the DLS of FXS model mice. Using high-frequency stimulation (HFS) to induce long-term depression (LTD) in acute striatal slices, we found that *Fmr1* KO mice exhibited significantly enhanced striatal LTD compared to WT littermates (**Fig. 1i, j**). These findings indicate that long-lasting synaptic plasticity is altered in the DLS of FXS model mice.

Next, we determined whether the synaptic alterations and RRBs observed in *Fmr1* KO mice result from exaggerated cap-translation in the dorsolateral striatum (DLS). First, we used surface sensing of translation (SUnSET) to label newly synthesized proteins in striatal coronal slices of *Fmr1* KO and WT mice. We observed a significant increase in *de novo* translation in the DLS of *Fmr1* KO mice compared to their WT littermates (**Fig. 2a**). Then, we investigated whether the enhanced protein synthesis exhibited by *Fmr1* KO mice in the DLS results from an increase in the interaction between the cap-binding translation initiation factor eIF4E and the initiation factor eIF4G. To test this hypothesis, we used a pull-down assay where m^7^GTP beads were incubated with DLS lysates from *Fmr1* KO and WT mice and found an increased eIF4G/eIF4E ratio in the *Fmr1* KO mice (**Fig. 2b**).

**Figure 2.**
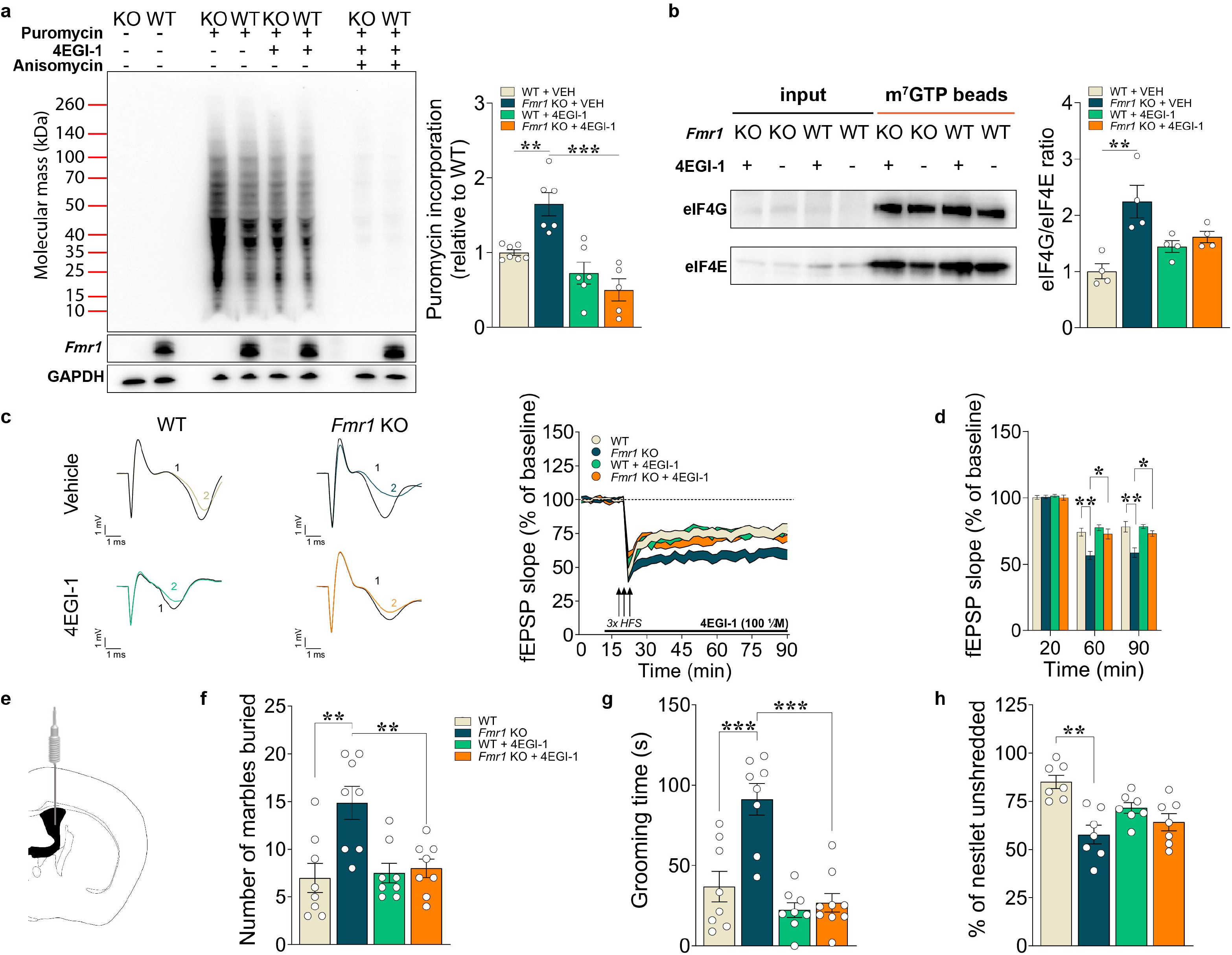
Increased *de novo* cap-dependent translation, cortico-striatal synaptic plasticity and repetitive/perseverative behavior exhibited by *Fmr1* KO mice are normalized by administration of 4EGI-1. *Fmr1* KO mice exhibit net increased *de novo* translation (**a**) and enhanced eIF4E– eIF4G interactions (**b**) in dorso-lateral striatum compared with controls. 4EGI-1 normalizes the exaggerated cap-dependent translation and decreases the enhanced eIF4E–eIF4G interactions in *Fmr1* KO mice. **a**, Representative western blots (left panel) and quantification of newly synthesized brain proteins in dorso-lateral striatum slices of *Fmr1* KO and WT mice, labelled with puromycin using the SUnSET method (see Methods). Summary plot (right panel) of puromycilation indicating increased *de novo* translation in dorso-lateral striatum of *Fmr1* KO mice compared to control and the effect of 4EGI-1 treatment (two-way ANOVA with Bonferroni’s multiple comparisons test; genotype x treatment interaction: *F_(1, 20)_* = 12.18, P=0.0023; genotype: *F_(1, 20)_* = 2.854, P=0.1067; treatment: *F_(1, 20)_* = 32.24, P<0.0001; **P < 0.01, *Fmr1* KO + VEH vs WT + VEH; ***P < 0.001, *Fmr1* KO + 4EGI-1 vs *Fmr1* KO + VEH; n= 5-7 independent lysates from 5-7 mice per group). **b**, Representative western blots (left panel) and quantification of pull-down assays with m^7^GTP beads (right panel) performed on lysates of dorsolateral striatal slices from *Fmr1* KO or WT mice and the respective treatment with 4EGI-1 or VEH (two-way ANOVA with Bonferroni’s multiple comparisons test; genotype x treatment interaction: *F_(1, 12)_* = 9.260, P=0. 0102; genotype: *F_(1, 12)_* = 16.10, P=0.0017; treatment: *F_(1, 12)_* = 0.2815, P=0.6054; **P < 0.01, Fmr1 KO + VEH vs WT + VEH; n= 4 independent lysates from 4 mice per group). Data are presented as mean□±□s.e.m. 4EGI-1 corrects enhanced cortico-striatal LTD shown by *Fmr1* KO mice without affecting LTD in WT mice (**c-d**). Cortico-striatal LTD was evoked by 3 trains of high frequency stimulation (HFS) and slices were treated with either 4EGI-1 (100 μM) or VEH applied 10 min before the tetanus and perfused for 70□min after tetanus. **c**, Representative field potentials before (1) and 70□min after tetanus (2) for different groups of slices are shown (left) and plot showing normalized fEPSP mean slope (± s.e.m. displayed every 2□min) recorded from coronal striatal slices from *Fmr1* KO and WT mice (right panel). **d**, Mean fEPSPs at baseline (20 min), at 60 (40 min after tetanus) and at 90 min (70 min after tetanus). 4EGI-1 normalizes fEPSP slope in *Fmr1* KO at both 60 and 90 min without affecting WT (two-way RM ANOVA, followed by Bonferroni’s multiple comparisons test; time x treatment interaction: *F_(6,136)_* = 4.672, *P*=0.0002; n=18 slices per group from 9 mice/genotype). Data are shown as mean ± s.e.m., \**P*< 0.05 and \*\**P*< 0.01 *Fmr1* KO versus WT mice. Inhibition of eIF4E-eIF4G interactions by 4EGI-1 reverses exaggerated repetitive and perseverative behaviours shown by *Fmr1* KO mice in the marble burying (**f**), grooming (**g**) and nestlet shredding (**h**) tests. **e**, Schematic for ICV injection of 4EGI-1. **f**, Treatment of *Fmr1* KO mice with 4EGI-1 (20μM) reduces the marble-burying behaviour. Summary plot of number of marbles buried after the 30 min marble burying test in *Fmr1* KO and WT mice treated with either 4EGI-1 or VEH (two-way ANOVA followed by Bonferroni post-hoc test was conducted; genotype x treatment interaction: *F_(1, 28)_* = 7.45, \**P*<0,05; genotype: *F_(1, 28)_* = 9.61, \*\**P*<0.01; treatment: *F_(1, 28)_*= 5.57, \**P*<0.05; *n*=8 mice/genotype/treatment). **g**, Summary plot of time spent grooming in the last 10 minutes interval of a 60-minute test (two-way ANOVA followed by Bonferroni post-hoc test was conducted; genotype x treatment interaction: *F_(1, 29)_* = 10.43, \*\**P*<0,01; genotype: *F_(1, 29)_* = 14.64, \*\*\**P*<0.001; treatment: *F_(1, 29)_*= 26.30, \*\*\**P*<0.001; *n*=8-9 mice/genotype/treatment). **h**, 4EGI-1 normalizes the enhanced repetitive/perseverative behaviour during the nestlet shredding test in *Fmr1* KO mice. Summary plot of percentage of unshredded nestlet during the nestlet shredding test (two-way ANOVA followed by Bonferroni post-hoc test was conducted; genotype x treatment interaction: *F_(1, 24)_* = 6.36, \**P*<0,05; genotype: *F_(1, 24)_* = 19.32, \*\*\**P*<0.001; treatment: *F_(1, 24)_*= 0.81, *P*=0.38; *n*=7 mice/genotype/treatment). All data are shown as mean ± s.e.m.; \**P*< 0.05, \*\**P*< 0.01, \*\*\**P*< 0.001 *Fmr1* KO *versus* WT mice.

Given our findings that the loss of FMRP results in a net increase in *de novo* translation (**Fig. 2a**) likely via enhanced eIF4E–eIF4G associations (**Fig. 2b**) in the DLS, we asked whether inhibiting the binding of eIF4E with eIF4G would rescue the exaggerated net protein synthesis (**Fig. 2a**), altered synaptic function (**Fig. 1**) and the locomotor and repetitive behavioral phenotypes exhibit by *Fmr1* KO mice (**Fig. 1f-h**). 4EGI-1, which prevents eIF4E–eIF4G interactions, has been reported to successfully rescue multiple ASD-like phenotypes in 4E-BP2 knockout and eIF4E transgenic mice^33,34^, as well as hippocampus-dependent memory impairments in FXS model mice^30^. We found that bath application of 4EGI-1 to cortico-striatal coronal slices from *Fmr1* KO mice normalized increased protein synthesis (**Fig. 2a**) and enhanced striatal LTD (**Fig. 2c,d**), indicating that those phenotypes are direct consequences of the increased association of eIF4E to eIF4G (**Fig. 2b**). Consistent with these observations, intracerebroventricular (ICV) injection of 4EGI-1^34^ normalized the RRBs exhibited by *Fmr1* KO mice without affecting WT controls (**Fig. 2f-h**). After 4EGI-1 administration, *Fmr1* KO mice exhibited a decrease in RRBs during the marble-burying task (**Fig. 2f**) and a reduction in the time spent grooming (**Fig. 2g**). Moreover, infusions of 4EGI-1 reduced the excessive proclivity in shredding the nestlet showed by *Fmr1* KO mice (**Fig. 2h**). Taken together, these findings support our hypothesis that the enhanced cap-dependent translation in the DLS occurs via increased eIF4E-eIF4G interactions and underlies morphological, synaptic, and RRBs displayed by FXS model mice.

Next, we asked whether the exaggerated cap-dependent translation observed in DLS of *Fmr1* KO mice is cell type-specific. We took advantage of fluorescent non-canonical amino acid tagging (FUNCAT), which allows visualization of newly synthesized proteins by measuring incorporation of a methionine analog (AHA) into nascent polypeptides. To investigate *de novo* translation specifically in Drd1- and Drd2-SPNs we generated two separate double mutant mouse lines to target and visualize direct and indirect pathway SPNs in living slice preparations, by crossing *Fmr1* KO mice with either Drd2-EGFP^+/−^ or Drd1a-tdTomato^+/−^ BAC transgenic mouse lines^35,36^. In cortico-striatal slices from the *Fmr1* KO/Drd2-EGFP^+/−^ and Drd1a-tdTomato^+/−^ BAC transgenic mice and their WT littermates, we observed a significant increase in newly synthesized proteins in Drd1-SPNs of *Fmr1* KO mice compared to WT controls (**Fig. 3**). In contrast, we did not observe a significant increase in *Fmr1* KO Drd2-SPNs (**Supplementary Fig. 1**). These findings indicate that the increase in *de novo* translation in the DLS of *Fmr1* KO mice occurs predominantly in the Drd1-SPNs.

**Figure 3.**
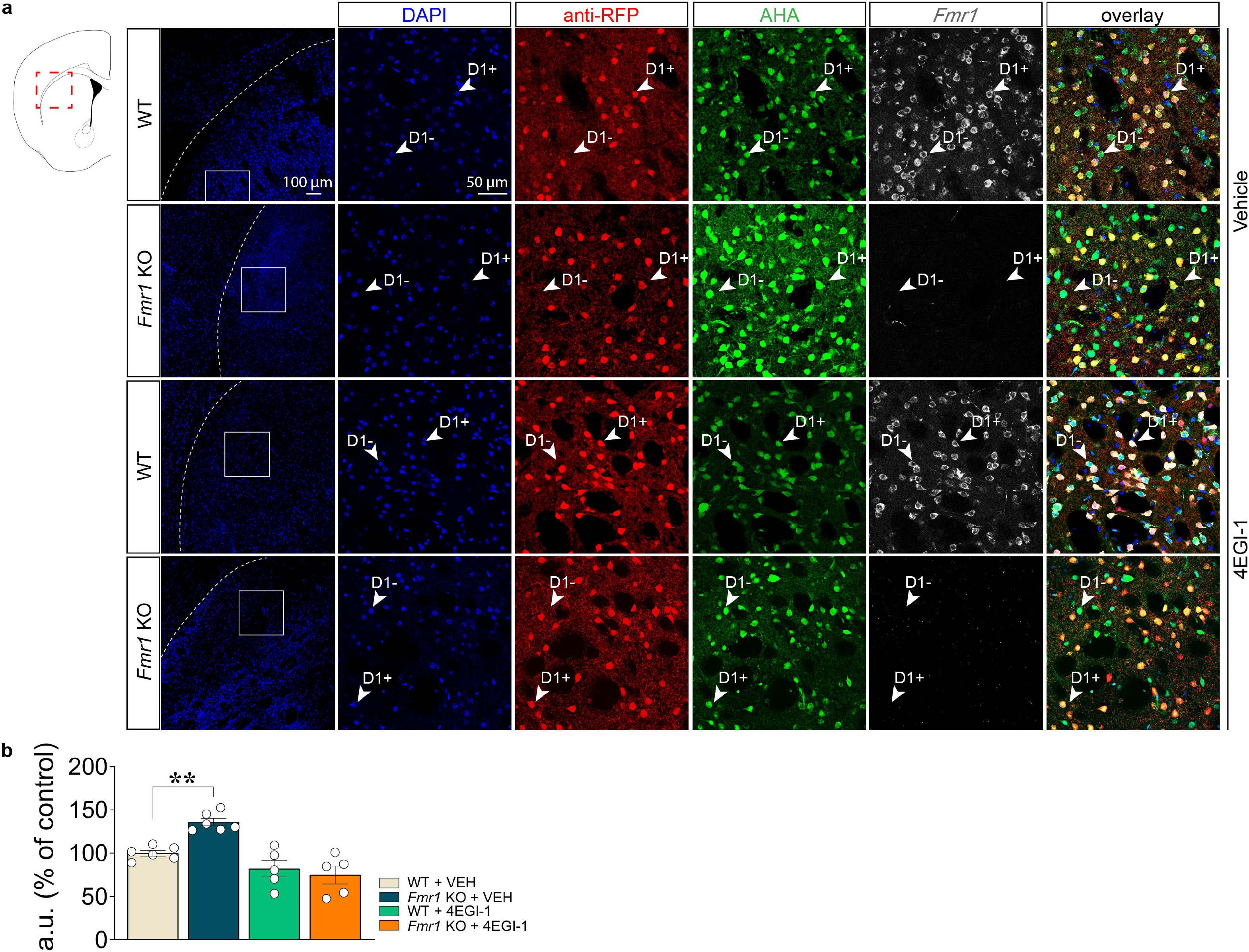
Lack of *Fmr1* results in dysregulated *de novo* translation in Drd1-SPNs, which is reversed by 4EGI-1. **a**, Representative DLS immunofluorescence images of DAPI (blue), anti-RFP (red), anti-FMRP (grey) and incorporation of AHA (green) detected by FUNCAT with alkyne-Alexa 488 in cortico-striatal slices from *Fmr1* KO/Drd1a-tdTomato BAC transgenic mice and their WT littermates (scale bar represents 50 μm) treated with VEH (first two rows from the top) or 4EGI-1(last two rows from the top). Insert in the first column is representative of the area magnified in each respective row (scale bar represents 100 μm). **b**, Quantification of increased AHA-alkyne-Alexa 488 signal in fluorescent arbitrary units (a.u.) expressed as % of control in Drd1-SPNs (anti-RFP+ neurons; red) from DLS *Fmr1* KO/Drd1a-tdTomato BAC transgenic mice and their WT littermates. Cell soma intensity was measured in ImageJ software (FIJI). Statistical significance was determined by using two-way ANOVA followed by Bonferroni’s multiple comparisons test (genotype x treatment interaction: *F_(1,18)_*= 9.35, \*\**P*<0,01; genotype: *F_(1,18)_*= 4.09, \**P*=0.05; treatment: *F_(1,18)_*= 30.95, \*\*\**P*<0.001). Data are shown as mean□±□s.e.m. of n□=□5/6 mice per group (average of n□=□20 somas per slice, n□=□2 slices per mouse, from three independent experiments) **P□<□0.01.

We hypothesized that the enhanced LTD in *Fmr1* KO mice compromises the balance of activity between the two populations of SPNs, leading to the RRBs and increased locomotor activity. Because long-term plasticity at cortico-striatal synapses can be differentially regulated and induced in direct (Drd1+) and indirect (Drd2+) pathway SPNs (dSPNs, iSPNs)^37^, we next investigated whether each pathway displayed abnormal synaptic connectivity in *Fmr1* KO mice compared to controls. First, we performed *in vivo* two-photon imaging through a window chronically implanted over the DLS of WT and *Fmr1* KO mice expressing tdTomato in dSPNs, while freely moving on a circular treadmill^38^. The Ca^2+^ indicator GCaMP6f^39^ was virally expressed in neurons to enable monitoring of Ca^2+^ dynamics in tdTomato-positive dSPNs and tdTomato-negative putative iSPNs^38^. Self-initiated forward locomotion was comparable between WT and *Fmr1* KO mice (**Supplementary Fig. 2**; mean velocity: *P* = 0.63; bout duration: *P* = 0.38; peak velocity: *P* = 0.40; total distance travelled: *P* =0.38; Mann-Whitney test, imaging sessions from 4 mice per genotype). The mean amplitude and frequency of Ca^2+^ transients imaged per active dSPN and iSPN did not differ across genotypes during locomotion, resulting in no net imbalance between pathways (**Supplementary Fig. 3**; Frequency: dSPN, *P* = 0.53; iSPNs, *P* = 0.53; pathway bias index, *P* = 0.32; Mann-Whitney test, n = 7 fields of view (FOVs) per genotype; Amplitude: dSPN, *P* = 0.46; iSPNs, *P* = 0.38; pathway bias index, *P* = 0.21; Mann-Whitney test, *n* = 7 FOVs per genotype). The total fraction of all imaged dSPNs and iSPNs recruited during self-initiated forward locomotion was also comparable between genotypes (**Supplementary Fig. 3**; dSPN, *P* = 0.15; iSPNs, *P* = 0.53; pathway bias index, *P* = 0.09; Mann-Whitney test, *n* = 7 FOVs per genotype).

Although these results do not, at face value, support a role for striatal dysfunction, the absence of behavioral phenotype in these mice led us to question whether the imaging approach, which requires the removal of a large area of the somatosensory cortex might disrupt the cortico-striatal circuits that mediate the behavioral phenotype of *Fmr1* KO mice (**Fig.1**). We therefore adopted a different approach to investigate SPNs activity. We subsequently recorded glutamatergic inputs in the form of mini excitatory postsynaptic currents (mEPSCs) from SPNs in DLS. Consistent with previous studies^35,36^, WT mice displayed mEPSCs of similar amplitude and frequency in dSPNs and iSPNs (**Fig. 4a-e**). However, in *Fmr1* KO mice mEPSCs were significantly increased in frequency, but not amplitude, in dSPNs compared to iSPNs (**Fig. 4a-e**). This selective upregulation suggests a potential imbalance in the excitatory drive between the direct and indirect pathways in *Fmr1* KO mice and increased frequency is likely due to increased synapse density or presynaptic release probability. We then assessed whether these functional changes were accompanied by structural alterations in SPNs. Accumulating evidence indicates that spine anomalies in both FXS individuals^40,41^ and ASD rodent models are recurring features^30,33,34^, however, a limited number of studies have a focus on FMRP’s role in regulating synaptic structure outside of the hippocampus and cortex. To determine whether FMRP regulates spine density in the SPNs of the DLS, we acquired z-stack confocal images of dSPNs and iSPNs (see methods) in DLS in coronal slices of *Fmr1* KO mice their littermate controls. Then, we applied a deconvolution technique^42^ to resolve the SPNs dendrite images and determined spine density in d and iSPNs. *Fmr1* KO mice exhibited a significant increase in spine density in dSPNs compared to wild-type littermates (**Fig. 4f**). In contrast, no difference was observed in iSPNs (**Fig. 4g**) and in the overall spine density in DLS of *Fmr1* KO mice compared to controls (**Supplementary Fig. 1c**). Our results highlight an important role for FMRP in controlling the number of striatal synapses specifically in the dSPNs and are largely consistent with previous reports from cortex and hippocampus showing FMRP as a key player in the regulation of synaptic structure and plasticity^21,22^.

**Figure 4.**
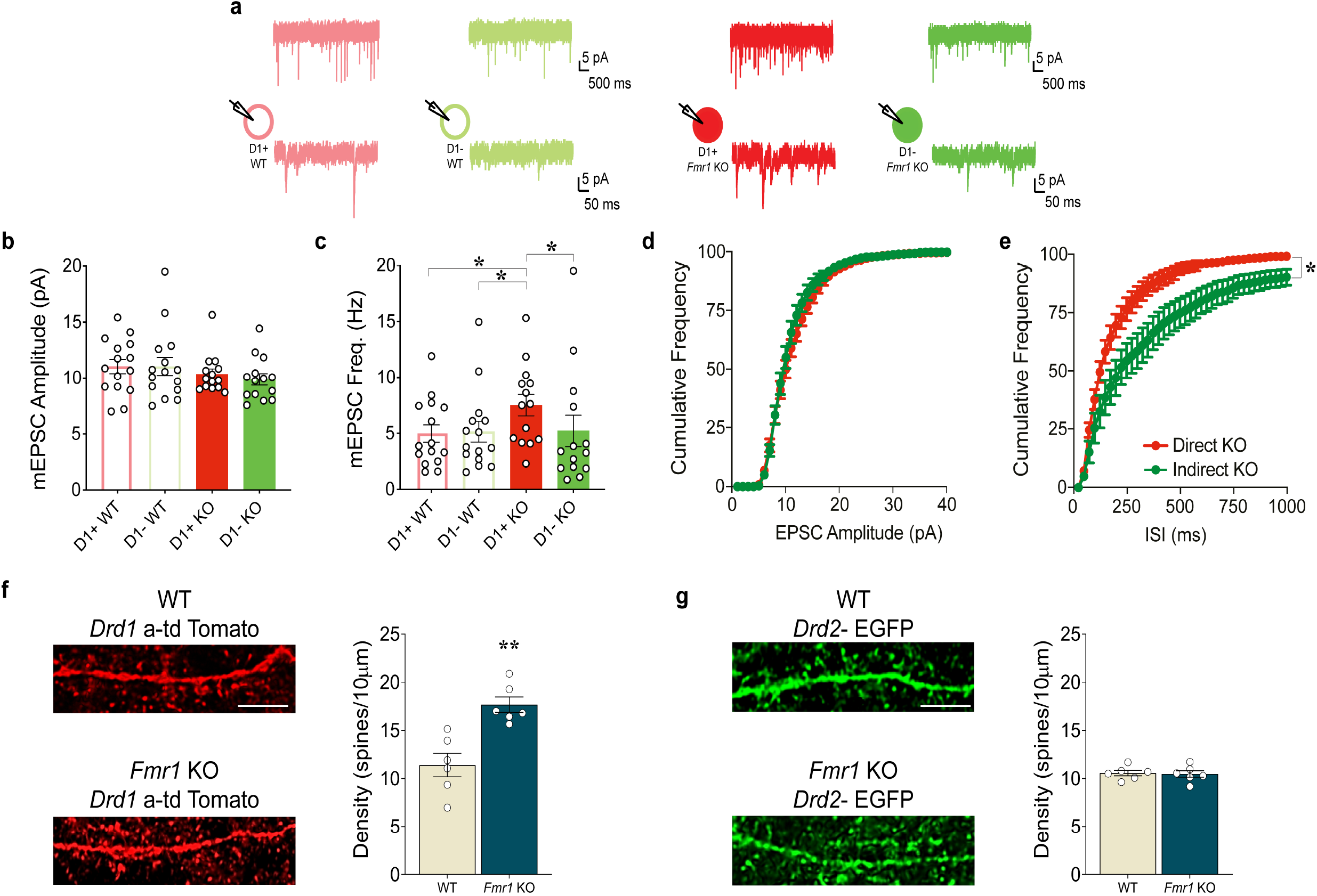
*Fmr1* KO mice exhibit altered synaptic function and spine density in dorsolateral striatum. *Fmr1* KO mice show increased mEPSC frequency but not mIPSC amplitude at Drd1-SPNs (**a-e**). **a**, Summary of mEPSC amplitude at Drd1- and Drd2-SPNs in control and *Fmr1* KO mice (no significant effects, Mann-Whitney test, mEPSC Amplitude WT Drd1- vs *Fmr1* KO Drd1- *P*=0.2898; WT Drd1+ vs *Fmr1* KO Drd1+ *P*=0.3048; *Fmr1* KO Drd1+ vs *Fmr1* KO Drd1- *P* =0.4013). **b**, Summary of mEPSC frequency at Drd1- and Drd2-SPNs in control and *Fmr1* KO mice. *Fmr1* KO exhibit increased mEPSC frequency at Drd1-SPNs (Mann-Whitney test, mEPSC Frequency WT Drd1+ vs WT Drd1- *P*=0.0910; WT Drd1- vs *Fmr1* KO Drd1- *P* =0.4706; WT Drd1+ vs *Fmr1* KO D1+ *P* =0.0259; *Fmr1* KO Drd1+ vs *Fmr1* KO Drd1- *P* =0.0186; *Fmr1* KO Drd1+ vs WT Drd1- *P* =0.041; *n* = 15 WT mice, *n* =14 *Fmr1* KO mice). Cumulative frequency plot of mEPSC amplitude (**c**) or ISI (**d**) recorded from Drd1- or Drd2-SPNs DL striatum of *Fmr1* KO mice (Kolmogorov-Smirnov test; e, amplitude *Fmr1* KO Drd1 + vs *Fmr1* KO Drd1- *P*=0.9206; f, ISI *Fmr1* KO Drd1+ vs *Fmr1* KO Drd1- *P*<0.0001). **e**, mEPSCs from neighboring Drd1- and Drd2-SPNs in WT control (left) or *Fmr1* KO (right) mice. Scale bar = 5 pA, 500 ms (top) or 5 pA, 50 ms (bottom). *Fmr1* KO mice exhibit increased dendritic spine density in dSPNs (**f,g**). High-magnification images (top) and quantification (bottom) of Drd1-SPNs (**f**) and Drd2-SPNs (**g**) spiny dendrites of WT and *Fmr1* KO mice (f, Mann-Whitney test, *P* < 0.01; g, Mann-Whitney test, *P* >0.99). Data are shown as mean ± s.e.m. of *n* = 6 mice/genotype (average of n = 3 slices per mouse; scale bar: 5 μm). kashi

### Trap-Seq of Drd1-SPNs reveals a coherent reduction in Rgs4

We next sought to determine the identity of mRNAs with altered translation in Drd1-SPNs. We employed translating ribosome affinity purification (Trap-Seq) that allows for cell-type-specific isolation of translating mRNAs using bacterial artificial chromosome (BAC) transgenic mouse lines engineered to express a GFP-tagged L10a ribosomal subunit in select cell populations^43^. Toward this end, we genetically expressed EGFP-tagged ribosomes in Drd1-SPNs by using a BAC transgenic Drd1-TRAP mouse line that shows a Drd1-SPNs-specific expression of EGFP-L10a within the striatum. Confocal imaging of coronal brain sections from Drd1-TRAP mouse confirmed expression of EGFP-L10a in Drd1-SPNs within the striatum (**Supplementary Fig. 4**). We then used anti-GFP antibodies on striatal lysates to immunoprecipitate the EGFP-L10a-labeled ribosomes and sequenced the co-purified mRNA. To decouple translational changes from alterations in total RNA expression, we also carried out RNA-Seq on RNA isolated from the whole striatal lysate (total).

Examination of the sequencing reads co-purifying with GFP-tagged ribosomes revealed an enrichment in known markers of striatal Drd1-SPNs and depletion of markers of Drd2-SPNs (**Supplementary Fig. 4a**). Ribosome association of the mRNAs coding for dopamine receptor D1 (*Drd1*), substance P (*Tac1*), and dynorphin (*Pdyn*) was enriched in the IP, compared to their RNA expression in the total lysates (**Supplementary Fig. 4a**). Dopamine receptor 2 (*Drd2*), adenosine 2a receptor (*Adora2a*), and enkephalin (*Penk*), which are characteristic markers of Drd2-SPNs, exhibited decreased ribosome association compared to their overall RNA expression in the striatum (**Supplementary Fig. 4a**).

Differential expression analysis of the immunoprecipitated RNA counts revealed 120 mRNAs (FDR < 0.1) with altered ribosome association in Drd1-SPNs of FXS model mice. Of the 120 mRNAs, 100 (83%) showed reduced ribosome association in FXS (**Fig. 5a-b**). Examination of mRNA abundances in whole striatal lysates, meanwhile, revealed 43 mRNAs that exhibited significant alterations in RNA expression in the striata of FXS mice (**Fig. 5a,c**). Similar to the pattern of alterations observed in ribosome-associated mRNA in dSPNs, 34 (79%) of the 43 mRNAs were down-regulated in FXS striata. The alterations observed in ribosome-associated mRNAs from Drd1-SPNs in FXS mice showed a moderate correlation with those observed in overall mRNA expression in FXS striata (Pearson’ r = 0.337; **Fig. 5d**). Moreover, the genes exhibiting significant RNA expression alterations in the whole striatum overlapped significantly with those exhibiting changes in ribosome association in dSPNs (n=13, excluding *Fmr1*; p = 2.8X10^−16^), with all genes showing downregulation in both assays (**Fig. 5d**). Consistent with these observations, gene set enrichment analysis (GSEA) revealed a reduction in mRNAs with gene ontologies (GOs) associated with cell adhesion, the synapse, and glutamate receptor signaling, and elevation in GOs associated with mitochondria and ribosomes in both the IP and total fractions in FXS mice (**Fig. 5f**, **Supplementary Fig. 4f**). Given that only ~37% of the cells in the striatum are dSPNs^44^, these results suggest that some of the changes in ribosome association in Drd1-SPNs may be driven by alterations in their RNA expression in FXS.

**Figure 5.**
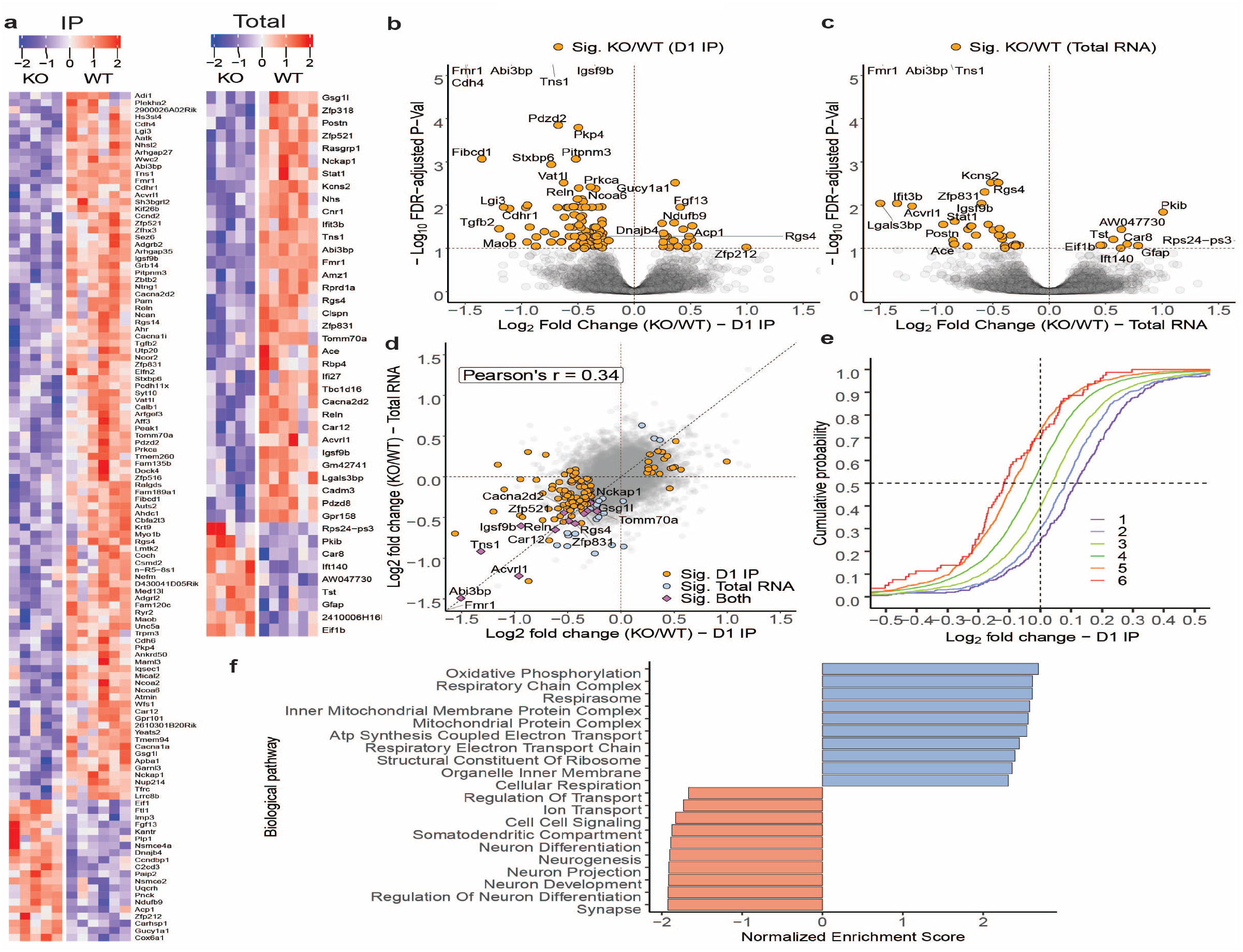
Translational profiling of Drd1 neurons in the striata of *Fmr1* KO mice. **a**, Heatmaps depicting ribosome associated mRNA expression (n=120) from dopamine Drd1 receptor-expressing medium spiny neurons (left; n=120) and overall striatal mRNA expression (n=43) of significantly different genes (FDR adjusted p-value< 0.1) between FXS and WT mice in translating ribosome affinity purification (left; IP) and RNA-seq (right; Total) assays. Each row in the heatmaps plots log2 transformed, centered and scaled (“row-normalized”) counts per million (CPM) values for significantly different genes. Majority of significantly different mRNAs show reduced ribosome association in D1 neurons (n=100↓, 20↑; IP) and reduced striatal mRNA expression (n=34↓, 9↑; Total) in FXS. **b**, Significance (FDR adjusted p-value) vs. log2-fold change (LFC) in ribosome association (IP) in D1 medium spiny neurons and **c**, overall striatal mRNA expression (Total) between FXS and WT mice. Messenger RNAs with significantly altered expression (FDR adjusted p-value< 0.1) in their respective assays, and absolute log2-fold-changes larger than 0.5 are labeled. **d**, Comparison of log-fold-changes (FXS/WT) in ribosome association in D1 MSNs (IP) against alterations in overall mRNA expression in the striata of FXS mice (Pearson’ r = 0.337). mRNAs exhibiting significant alterations in both D1 neuron ribosome association and overall RNA expression are labeled. **e**, Cumulative distribution of log2-fold-changes (FXS/WT) in ribosome association in D1 MSNs (IP) in the striatum of FXS model mice, as a function of coding sequence length. Messenger RNAs are divided into six bins in ascending order of their CDS lengths, with bin 1 harboring mRNAs with the shortest CDSs, and bin 6 the longest. mRNAs with the longest CDSs are particularly likely to exhibit reduced ribosome association in D1 MSNs of FXS, while those with the short CDSs are especially likely to show increased ribosome association. A gradation of effects can be observed between these two extremes. **f**, Top 10 gene ontologies exhibiting alterations in ribosome association in D1 MSNs of FXS mice. RNA co-purifying with ribosomes from D1 MSNs of FXS mice is enriched in transcripts coding for mitochondrial processes and ribosomal proteins, and shows reductions in GOs associated with cell adhesion, glutamate receptor signaling, and the synapse.

Earlier work has suggested that the coding sequence (CDS) length of an mRNA is associated with alterations in ribosome association in FXS^31,45–47^. To examine whether the CDS length of an mRNA dictates its ribosome association in dSPNs of FXS model mice, we ordered all mRNAs ascendingly by their CDS lengths, divided them into six color-coded bins, and evaluated their log2-fold changes against their FDR-adjusted p-values. Consistent with the prior observations, mRNAs with the longest CDSs were enriched in genes with significantly reduced ribosome association in dSPNs of mice lacking FMRP, while those with the shortest CDSs were enriched in genes with significantly increased ribosome association (**Supplementary Fig. 4b**). Evaluation of RNA expression changes in the striatum in FXS mice revealed a slight trend towards the same pattern observed in ribosome association in dSPNs (**Supplementary Fig. 4c**). Examination of the cumulative distribution of the alterations in each CDS length bin revealed a positive-to-negative gradation of log2-fold-changes in dSPNs of FXS model mice (**Fig. 5e**). Over 75% of mRNAs with the longest CDSs exhibited reduced ribosome association, whereas fewer than 30% of those with the shortest CDSs were altered in the same direction. A gradual trend in the direction of alteration was observed between these two extremes. Bin-wise examination of RNA expression in the striata of FXS mice showed that alterations in mRNAs with short CDSs closely tracked those in ribosome association in dSPNs. (**Supplementary Fig. 4d**).

To confirm that our results were not simply the consequence of the binning thresholds we employed, and to directly compare the CDS length-dependent alterations in IP and striatal mRNA expression (total) in FXS, we ordered mRNAs into 50 bins by their CDS length, with each bin harboring the same number of mRNAs. Evaluating the average LFCs in ribosome association in dSPNs of FXS mice across the 50 CDS length bins confirmed the positive-to-negative linear trend in alterations with CDS length (**Supplementary Fig. 4e**). The average LFCs in whole striatal RNA expression of mRNAs with short CDSs closely tracked the LFCs in ribosome association in dSPNs of FXS mice. For mRNAs with longer CDSs, the average LFCs in RNA expression were predominantly negative, but lay closer to zero and diverged from the highly negative average LFCs observed in ribosome association in dSPNs of mice lacking FMRP.

### M4R positive allosteric modulator VU0152100 corrects enhanced cortico-striatal LTD and RRBs in *Fmr1* KO mice

Notably, TRAP-Seq of dSPNs revealed significant reduction of *Rgs4* in RNA that co-purified with EGFP-tagged ribosomes derived form Drd1-SPNs – as well as in total striatal RNA – of Fmr1 KO mice (**Fig. 5b,c,d**). Within the striatum, G-protein signaling (RGS) 4 GTPase accelerating enzyme, interacts with different receptor systems, including the muscarinic 4 receptor (M4R)^48,49^, and is necessary for plasticity, specifically dopamine-mediated regulation of LTD in dorsal striatum^50^. Among those transcripts that were significantly decreased in RNA bound to ribosomes in dSPNs of *Fmr1* KO mice, we found the. M4R is a Gi/o protein-coupled receptor and its activity is mediated by the inhibition of adenyl cyclase (AC), which reduces cyclic adenosine monophosphate (cAMP) levels and close voltage-gated Ca^2+^ channels (VGCCs)^51^. A previous study on the FXS mouse model showed that inhibition of M4R results in a significant increase in protein synthesis in hippocampal slices of both WT and *Fmr1* KO mice, whereas VU0152100, an M4R positive allosteric modulator (PAM) has been shown to normalize, exaggerated hippocampal protein synthesis and mGluR-LTD in FXS model mice^52^.

To gain insight into the role of Rgs4 within dSPNs underlying the expression of RRBs, we prepared cortico-striatal coronal slices from *Fmr1* KO and WT mice for cortico-striatal LTD studies with bath application of 5 μM VU0152100, a concentration which was shown to enhance M4R function, reduce protein synthesis and normalize mGluR-LTD in hippocampal slices of *Fmr1* KO mice^52^. We found that VU0152100 normalized enhanced cortico-striatal LTD exhibited by *Fmr1* KO mice without affecting LTD in WT mice (**Fig. 6a,b**). Next, we sought to determine the impact of of M4R signaling modulation on RRBs. *Fmr1* KO and WT mice were treated with VU0152100 (56 mg/kg; i.p.) and then tested for RRBs. M4R PAM normalized the aberrant RRBs exhibited by *Fmr1* KO mice without affecting their WT littermates (**Fig. 6c-e**). After VU0152100 administration, *Fmr1* KO mice exhibited reduced RRBs in the marble-burying task (**Fig. 6c**) and engaged less in grooming activity (**Fig. 6d**). Finally, the VU0152100 treatment reduced the excessive proclivity in nestlet shredding displayed by the *Fmr1* KO mice (**Fig. 6e**). Taken together, these results indicate that enhancing M4R signaling, possibly via Rgs4, corrects exaggerated cortico-striatal LTD and the RRBs exhibited by *Fmr1* KO mice.

**Figure 6.**
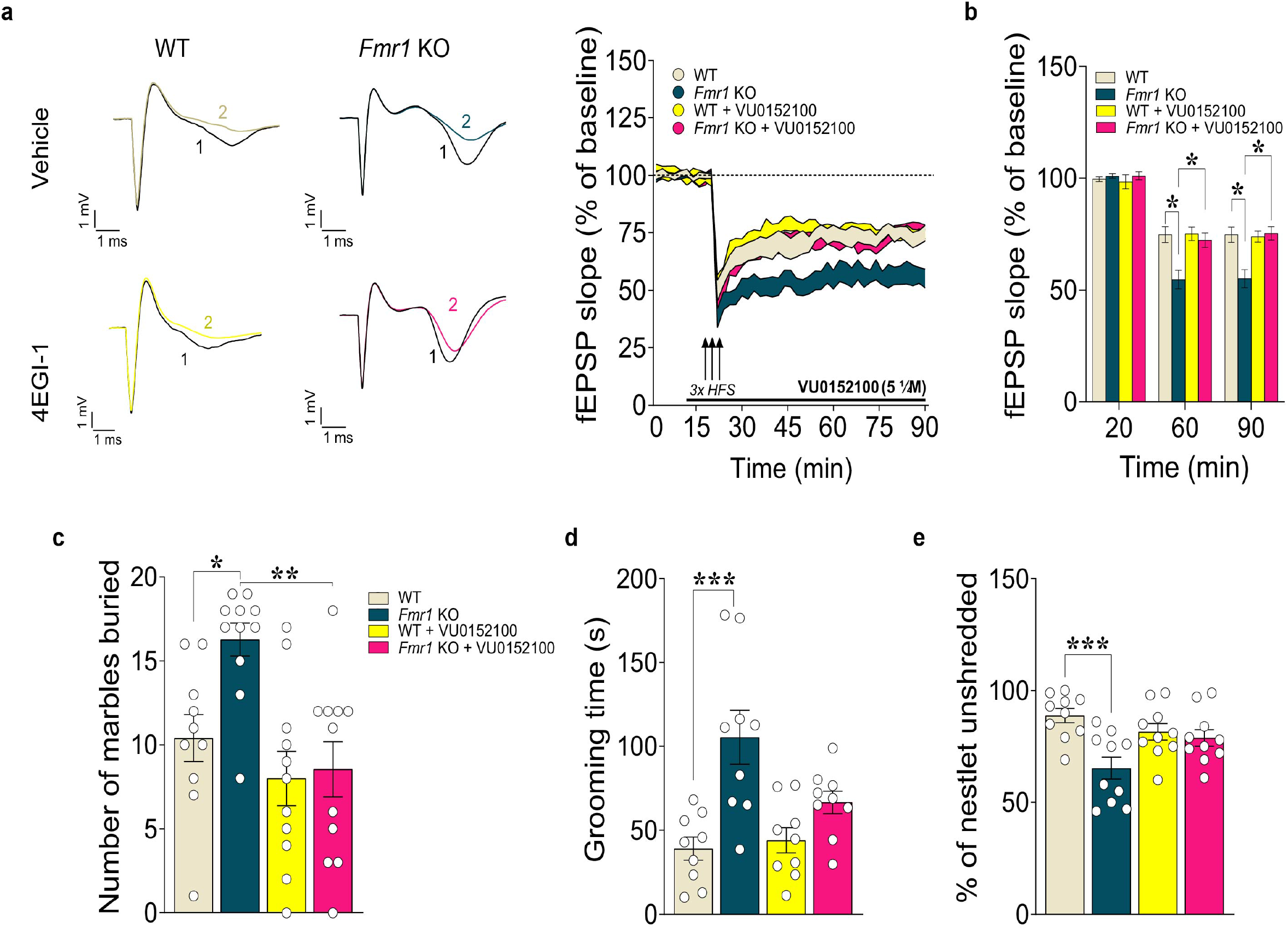
VU0152100 corrects excessive repetitive behavior and exaggerated cortico-striatal LTD in *Fmr1* KO mice. VU0152100 normalizes enhanced cortico-striatal LTD shown by *Fmr1* KO mice without affecting LTD in WT mice (**a-b**). Cortico-striatal LTD was evoked by 3 trains of high frequency stimulation (HFS) and slices were treated with either VU0152100 (5 μM) or VEH applied 10 min before the tetanus and perfused for 70□min after tetanus. **a**, Representative field potentials before (1) and 70□min after tetanus (2) for different groups of slices are shown (left) and plot showing normalized fEPSP mean slope (± s.e.m. displayed every 2□min) recorded from coronal striatal slices from *Fmr1* KO and WT mice (right panel; *n*=14 slices from 7 mice/genotype/treatment). **b**, Mean fEPSPs at baseline (20 min), at 60 (40 min after tetanus) and at 90 min (70 min after tetanus). LTD was significantly enhanced in *Fmr1* KO striatal slices treated with VEH at both 60 min and 90 min compared with WT VEH, whereas VU0152100 normalizes fEPSP slope in *Fmr1* KO at both 60 and 90 min without affecting WT (two-way RM ANOVA, followed by Bonferroni’s multiple comparisons test; time x treatment interaction: *F_(6,104)_* = 4.44, \*\*\**P*<0.001; n=14 slices per group from 7 mice/genotype). Data are presented as mean□±□s.e.m., \**P*< 0.05 *Fmr1* KO *versus* WT mice. The M4R PAM VU0152100 reverses exaggerated repetitive and perseverative behaviours shown by *Fmr1* KO mice in the marble burying (**c**), grooming (**d**) and nestlet shredding (**e**) tests. **c**, Treatment of *Fmr1* KO mice with VU0152100 (56 mg/kg; i.p.) reduces the marble-burying behaviour. Summary plot of number of marbles buried after the 30 min marble burying test in *Fmr1* KO and WT mice treated with either VU0152100 or VEH (two-way ANOVA followed by Bonferroni’s post-hoc test was conducted; genotype x treatment interaction: *F_(1,39)_*= 3.42, *P*=0.07; genotype: *F_(1,39)_*= 4.96, \**P*<0.05; treatment: *F_(1,39)_*= 12.34, \*\**P*<0.01; *n*=10-11 mice/genotype/treatment). **d**, VU0152100 improves the exaggerated grooming in *Fmr1* KO mice. Summary plot of time spent grooming in the last 10 minutes interval of a 60-minute test (two-way ANOVA followed by Bonferroni’s post-hoc test was conducted; genotype x treatment interaction: *F_(1,32)_*= 4.67, \**P*<0.05; genotype: *F_(1,32)_*= 19.31, \*\*\**P*<0.001; treatment: *F_(1,32)_*= 2.79, *P*=0.10; *n*=9 mice/genotype/treatment). **e**, *Fmr1* KO shows reduced repetitive/perseverative behaviour after treatment with VU0152100 in the nestlet shredding test. Summary plot of percentage of unshredded nestlet during the nestlet shredding test (two-way ANOVA followed by Bonferroni’s post-hoc test was conducted; genotype x treatment interaction: *F_(1,36)_*= 6.99, \**P*<0.05; genotype: *F_(1,36)_*= 11.10, \*\**P*<0.01; treatment: *F_(1,36)_*= 0.62, *P*=0.43; *n*=10 mice/genotype/treatment). All data are shown as mean ± s.e.m.; \**P*< 0.05, \*\**P*< 0.01 *Fmr1* KO *versus* WT mice.

### Conditional deletion of *Fmr1* in dSPNs leads to a net increase in protein synthesis and excessive RRBs in mice

In order to evaluate the cell type-specific contribution of FMRP expression in Drd1-SPNs to RRBs in FXS, we generated mice containing Drd1 promoter-driven Cre transgene (GENSAT Project at Rockefeller University)^53^ and a conditional allele of *Fmr1* (*Fmr1*loxP; termed *Fmr1*^f/f^; **Fig. 7a**)^54^. The expression of the Cre transgene and the *Fmr1*loxP allele was determined using PCR-specific primers (**Fig. 7b**). The resulting conditional knockout mice (*Fmr1*^f/f^ Drd1-Cre), which lack FMRP expression in Drd1-SPNs and their littermate controls (Fmr1^+/+^ Drd1-Cre) were used to test the causal role of Drd1-SPNs specific deletion of *Fmr1* in RRBs expression.

**Figure 7.**
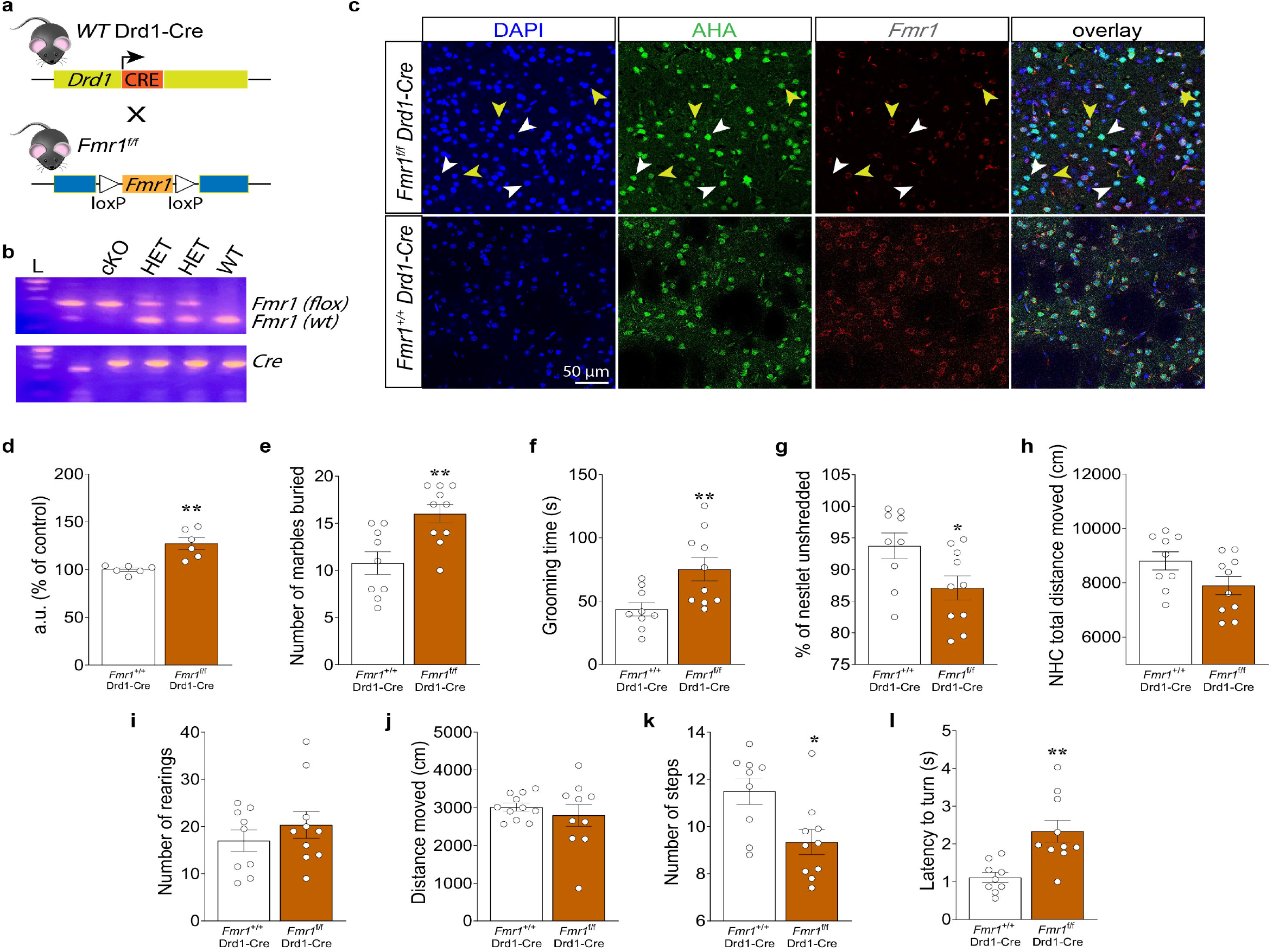
Selective deletion of *Fmr1* from Drd1-SPNs results in dysregulated *de novo* translation and excessive repetitive behavior in mice. **a**, Schematic representation of Drd1-neuron specific deletion of *Fmr1* in *Fmr1*^f/f^ mice crossed with Drd1-Cre mice. **b**, PCR identification of alleles of *Fmr1*loxP and Drd1-driven Cre. **c**, Representative DLS immunofluorescence images of DAPI (blue), anti-FMRP (red), and incorporation of AHA (green) detected by FUNCAT with alkyne-Alexa 488 in cortico-striatal slices from *Fmr1*^f/f^ Drd1-Cre and *Fmr1*^+/+^ Drd1-Cre mice (scale bar represents 50 μm). White arrows indicate Drd1-SPNs (green) and FMRP (red) co-staining; yellow arrows indicate non-Drd1-SPNs and FMRP (red) negative staining. **d**, Quantification of increased AHA-alkyne-Alexa 488 signal in fluorescent arbitrary units (a.u.) expressed as % of control in Drd1-SPNs from DLS of *Fmr1*^f/f^ Drd1-Cre vs. *Fmr1*^+/+^ Drd1-Cre mice. Cell soma intensity was measured in ImageJ software (FIJI). Statistical significance was determined by using Student’s t test (*Fmr1*^f/f^ Drd1-Cre vs. *Fmr1*^+/+^ Drd1-Cre mice; unpaired *t* test; t*_(10)_*□=□4.28, \*\**P*□<0.01. Data are shown as mean□±□s.e.m. of n□=□6 mice per group (average of n□=□20 somas per slice, n□=□2 slices per mouse, from three independent experiments). (**e-l**) *Fmr1*^f/f^ Drd1-Cre mice exhibited exaggerated repetitive and perseverative behaviors, but normal spontaneous locomotor activity compared to their *Fmr1*^+/+^ Drd1-Cre control mice. **e**, Summary plot of number of marbles buried after the 30 min test (unpaired *t* test, t*_(17)_*= 3.40, \*\**P* <0.01; *n*= 9-10 mice/genotype). **f**, Summary plot of time spent grooming in the last 10 minutes interval of a 60 minutes test (unpaired *t* test, t*_(17)_*= 2.90, \*\**P* <0.01; *n*= 9-10 mice/genotype). **g**, Summary plot of percentage of unshredded nestlet during the nestlet shredding test (unpaired *t* test, t*_(17)_*= 2.38, \**P* <0.05; *n*= 9-10 mice/genotype). **h**, Summary plot of the novelty-induced locomotor activity expressed as a novel home cage (NHC) distance moved (cm) in a 60 minutes test during novel home cage test (unpaired *t* test, t_(*17*)_= 1.92, *P*=0.072; *n*=9-10 mice/genotype). **i**, Summary plot of vertical activity expressed as number of rearing episodes (number of counts) during the cylinder test over 5 min (unpaired *t* test, t*_(17)_*= 0.91, *P*=0.37; *n*=9-10 mice/genotype). **j**, Summary plot of spontaneous locomotor activity expressed as distance moved (cm) during the open field test over 15 min (unpaired *t* test, t*_(19)_*= 0.75, *P*=0.47; *n*=10-11 mice/genotype). *Fmr1*^f/f^ Drd1-Cre showed motor impairment in motor tests specific for assessing striatal-based locomotor activity (**k-l**) when compared to their littermates control mice. **k**, Summary plot of average number of steps during drag test (unpaired *t* test, t*_(17)_*= 2.80, \**P* <0.05; *n*=9-10 mice/genotype). **l**, Summary plot of latency to turn (s) during pole test (unpaired *t* test, t*_(17)_*= 3.69, \*\**P* <0.01; *n*=9-10 mice/genotype). Mice were analyzed using Student’s *t* test. \**P* < 0.05, \*\**P* < 0.01 *Fmr1*^f/f^ Drd1-Cre mice different from age-matched *Fmr1*^+/+^ Drd1-Cre control mice. All data are shown as mean ± s.e.m.

Given our findings that the increase in *de novo* translation in the DLS of *Fmr1* KO mice (**Fig. 2**) is mostly attributable to dSPNs (**Fig. 3**; **Supplementary Fig. 1**), we investigated the effect of removing FMRP on *de novo* protein synthesis in Drd1-expressing cells, including SPNs (**Fig. 7c,d**). Cortico-striatal slices from *Fmr1*^f/f^ Drd1-Cre mice and their control littermates were subjected to FUNCAT to detect the rate of newly synthesized proteins. We observed a significant increase (~30%) in *de novo* translation in dSPNs in the DLS of *Fmr1*^f/f^ Drd1-Cre mice compared to wild-type Drd1-Cre littermates (**Fig. 7c,d**). To confirm that the increased *de novo* translation due to the selective ablation of FMRP in Drd1-expressing cells is sufficient to facilitate locomotor activity and drive the expression of RRBs as shown by *Fmr1* KO mice, we examined the conditional *Fmr1* KO mice and their littermate controls in a set of behavioral tests specific for motor abilities and RRBs. Consistent with the findings with the *Fmr1* KO mice (**Fig. 1f-h**), *Fmr1*^f/f^ Drd1-Cre mice displayed greater number of marbles buried compared to controls in the MB test (**Fig. 7e**). Likewise, we found that *Fmr1*^f/f^ Drd1-Cre mice engaged in significantly more grooming activity compared to controls (**Fig. 7f**) and shredded significantly more of their nestlets compared to *Fmr1*^+/+^ Drd1-Cre mice (**Fig. 7g**). However, selective deletion of *Fmr1* in Drd1-expressing cells did not affect spontaneous locomotor activity in *Fmr1*^f/f^ Drd1-Cre mice compared to their controls when tested for either the NHC (**Fig. 7h**), the cylinder (**Fig. 7i**) or the OF (**Fig. 7j**) test. Surprisingly, we found that *Fmr1*^f/f^ Drd1-Cre mice exhibited significantly reduced locomotor ability than controls in both drag test (**Fig. 7k**) and pole test (**Fig. 7l)** resulting in an opposite behavioral outcome compared to the *Fmr1* KO mice for both tests (**Fig. 1d,e**).

Collectively, these data support the idea that FMRP loss in dSPNs alters control of motor functions and is necessary and sufficient to trigger RRBs via the disruption of the FMRP-dependent translational control in SPNs. In contrast, selective deletion of *Fmr1* in Drd1-expressing cells is not sufficient to increase spontaneous locomotor activity.

## Discussion

In the present study, we sought to unveil the pathological synaptic and molecular mechanisms underlying RRBs and hyperactivity in FXS focusing on DLS SPNs. Most neurons lacking FMRP exhibit exaggerated protein synthesis, which contributes to the synaptic, structural, and behavioural deficits associated with FXS ^19,21,22^. Therefore, a number of therapeutic strategies have focused on reducing the enhanced translation to rescue FXS-associated phenotypes^24,30,32,55^. The properties of FMRP as a translation regulator have been intensely investigated in the cortex and hippocampus, leaving the striatum relatively unexplored. This is the first study showing that loss of FMRP in the DLS SPNs results in a net increase in cap-dependent translation, which contributes to altered synaptic plasticity, spine density, and RRBs in *Fmr1* KO mice. Our findings suggest that striatal spine morphology and synaptic plasticity rely on proper translational control, which is disrupted in SPNs lacking FMRP. Hence, *Fmr1* KO mice display aberrant motor behaviors that are likely protein synthesis-dependent and cortico-striatal in nature.

We found that the increase in DLS protein synthesis occurs via enhanced eIF4E-eIF4G interactions in the SPNs of *Fmr1* KO mice. Evidence suggests that gain of function mutations in eIF4E are associated with autistic behaviors^56^, suggesting a link between elevated cap-dependent translation and ASD. Studies utilizing genetic manipulation of proteins involved in cap-dependent translation^33,34,57,58^ further support our findings. Deletion of the eukaryotic translation initiation factor 4E-binding protein 2 (4E-BP2)^33^, an eIF4E repressor downstream of mTORC1, or the overexpression of eIF4E^34^ results in increased translation and ASD-like behaviors in mice, including hyperactivity and RRBs. eIF4E transgenic mice exhibit also enhanced cortico-striatal LTD compared to control and inhibition of the formation of the eIF4F complex rescues increased protein synthesis, enhanced LTD and RRBs in mice^34^. Consistent with these studies^30,33,34^, we found that increased binding of eIF4E with eIF4G may account for the net increase in protein synthesis in the DLS, which is responsible for the enhanced cortico-striatal LTD and, ultimately RRBs in *Fmr1* KO mice (**Fig. 3**). Notably, 4EGI-1 did not affect the expression of LTD in WT mice suggesting, that altered formation of eIF4E-eIF4G complex, and its activity is necessary for the expression of the enhanced cortico-striatal LTD in *Fmr1* KO mice but is not sufficient for this form of synaptic plasticity in WT mice. This is consistent with similar observations from mGluR-LTD experiments performed in presence of either cercosporamide^59^, an inhibitor of eIF4E phosphorylation, or 4EGI-1^30^ in the hippocampus of WT and FXS model mice.

To our knowledge, this is the first study showing that *Fmr1* KO mice exhibit functional changes at cortico-striatal synapses consistent with structural alterations in dSPNs population of the DLS (**Fig. 4**). HFS in DLS results in enhanced cortico-striatal LTD in *Fmr1* KO mice compared to WT, suggesting altered striatal information processing in FXS, which likely compromises the balanced activity of the two striatal pathways. mGluR-LTD is expressed throughout the striatum and contributes to balance the direct and indirect pathways, dominated by a long-lasting inhibitory effect on the indirect pathway^60^. In line with our results, eIF4E-transgenic mice, a model of ASD, exhibit enhanced cortico-striatal LTD and are unable to form new motor patterns and disengage from previously learned motor behaviours^34^. Also, mice with deletion of the synapse-associated protein 90/postsynaptic density protein 95-associated protein 3 (Sapap3), a synaptic protein that binds Shank3, display robust self-grooming behavior that was correlated with synapses elevated mGluR5 signaling and synaptic dysfunction at cortico-striatal synapses, and which was alleviated by mGluR5 inhibition^61,62^. Circuit-specific cortico-striatal functional connectivity imbalance has emerged as the main neural underpinning of RRBs in ASDs^9^ and that long-term plasticity at cortico-striatal synapses can be differentially regulated and induced in the two populations of SPNs^37^. Therefore, it is not surprising that examination of spontaneous mEPSCs in DLS revealed an increase in the frequency, but not the amplitude, of excitatory events exclusively in dSPNs of FXS mice (**Fig. 4**). Thus, our data strongly suggest that this selective upregulation of excitatory inputs toward the direct pathway may trigger the synaptic transmission imbalance within the cortico-striatal circuits responsible for the expression of RRBs and hyperactivity in FXS. Consistent with this idea, we observed a selective increase in the frequency of spontaneous mEPSCs in dSPNs (**Fig. 4c,e**), likely due to a selective increased synaptic density in Drd1-SPNs in FXS mice (**Fig. 4f,g; Supplementary Fig.1c**), which may reflect an enhanced number of synaptic contacts at cortico-striatal synapses in the DLS of FXS mice. In agreement with our findings, cell-type specific deletion of Tsc1 in dSPNs was shown to impair endocannabinoid-mediated long-term depression (eCB-LTD) at cortico-dSPN synapses enhancing corticostriatal synaptic drive and resulting in enhanced motor learning^63^. In addition, mice carrying neuroligin mutations exhibit repetitive behaviors associated with a selective decrease of synaptic inhibition onto dSPNs and striatal synaptic function in the nucleus accumbens (NAc)^64,65^. It is important to note that the analysis of the overall spine density in the DLS (dSPNs + iSPNs) of *Fmr1* KO mice show no difference compared to controls. In addition, a previous study using a cortical-striatal co-culture model of FXS reported reduced dendritic spine density in SPNs lacking FMRP^66^. However, the same group^67^, and others^68^, have reported increased SPNs spine density in the nucleus accumbens (NAc) core subregion of *Fmr1* KO mice and a reduction of mature spines^69^ in the DLS, suggesting that absence of FMRP *in vivo* drives different dendritic phenotypes, even within striatal subregions^66^. On the other hand, it has been postulated that different neuron and synapse types may adapt differently to the lack of FMRP^70^. Consistent with this notion, mGlu5-mediated endocannabinoid (eCB) activity at GABAergic synapses in the dorsal striatum of *Fmr1* KO mice is increased^71^, whereas eCB-LTD is abolished in the ventral striatum^70^.

The cortico-striatal functional changes we found are associated with structural alterations in the dSPNs of FXS mice and are consistent with the role of FMRP in regulating synaptic structure and plasticity reported in both cortex and hippocampus ^72,73^. Thus, hyperactivity and RRBs may result from enhanced activity within the basal ganglia circuitry arising from mechanisms that link dendritic spine pathology to circuit abnormalities relevant to atypical behavior^74^. Abnormalities in striatal structure and function have been observed across different preclinical models of ASD^3,30,33^ and have demonstrated that the composition of cortico-striatal synapses plays a key role in striatum-based ASD-like behaviors. Evidence from studies on ASD models^27^ indicates that hyperactivity is generally accompanied by RRBs, suggesting an impaired coherence across cortico-striatal circuits in the expression of both phenotypes^75^. For example, mice with mutations in the *SCN1A* gene exhibit ASD-like phenotypes, including hyperactivity, stereotypic self-grooming, and circling behaviors, along with increased cortical excitation^76^. *Fmr1* KO mice exhibit increased locomotor activity and excessive proclivity in engaging in RRBs, which are consistent with previous studies on FXS model mice^32,77^. In addition, we show for the first time that *Fmr1* KO mice exhibited greater motor facilitation in specific behavioral tests that assess striatum-driven locomotor activity.

Several lines of evidence from our study suggest that dSPNs are impacted by the loss of FMRP. FUNCAT experiments indicate that dSPNs exhibit a robust and significant increase in *de novo* translation in *Fmr1* KO mice and selective deletion of FMRP in Drd1-SPNs corroborated these findings. Fmr1^f/f^ Drd1-Cre mice exhibited a significant increase in *de novo* translation in Drd1-SPNs and exhibited RRBs to a similar extent as *Fmr1* KO mice, despite a quite opposite locomotor phenotype. FMRP loss in Drd1-expressing cells is therefore necessary and sufficient for the expression of RRBs but does not alter spontaneous locomotor activity. The DLS is poised to be a hub in the control of different locomotor abilities as it receives major input from dopaminergic nigral innervation, and cortical regions^78^. The selective deletion of *Fmr1* in Drd1-SPNs results in a disruption of the functional antagonism between the direct and indirect striatofugal pathways leading to an opposite motor phenotype in those mice. Therefore, loss of FMRP only in Drd1-SPNs may not be sufficient to trigger specific locomotor behaviors such as novelty-induced and/or spontaneous locomotor activity, suggesting that an increase in protein synthesis in either both dSPNs and iSPNs or cortical/dopaminergic neuronal inputs is required to generate a hyperkinetic phenotype in FXS model mice. In line with this notion, we found that 4EGI-1 ICV injection normalized the RRBs exhibited by *Fmr1* KO mice without affecting their WT controls, whereas hyperactivity was not rescued by the administration of 4EGI-1 (data not shown), suggesting a preferential role of dysregulated translation in the genesis of RRBs versus hyperactivity in FXS.

We attempted to determine whether pathological changes in Drd1-SPNs was due to the altered translation of specific mRNAs. TRAP-seq of Drd1-SPNs and RNA-seq of striatal lysates from *Fmr1* KO mice revealed a reduction in ribosome association and total expression of mRNAs associated with cell adhesion, the synapse, and voltage-gated calcium channels, and elevation in those associated with the ribosome and mitochondria. Aberrant expression of mitochondrial genes and subsequent changes in metabolic processes in the mitochondria have been observed in FXS model mice^79^. Similarly, genome-wide association studies have demonstrated that alterations in synaptic genes, including those encoding cell adhesion molecules and voltage-gated calcium channels, are particularly relevant to the pathogenesis of ASD^80^. Our results revealed that in the striatum of FXS model mice, alterations in expression of mRNAs with short CDSs closely track that of ribosome association in Drd1-SPNs (**Supplementary Fig. 4e**). This correspondence suggests that the elevation of LFCs in ribosome association observed in short CDS mRNAs may simply be due to an elevation in their RNA expression, rather than due to a fundamental change in translation. For longer CDS mRNAs, however, the LFCs in striatal RNA expression in FXS mice do not fully explain the large negative LFCs in ribosome association observed in Drd1-SPNs. The similarity of ribosome-bound mRNA and overall mRNA expression also was noted in a previous Trap-Seq study of hippocampal CA1 neurons in FXS mice^52^. Upon examination of the differentially expressed transcripts, we found an increase in the expression of eIF1 and other ribosome-associated mRNAs (e.g., Paip2) in *Fmr1* KO Drd1-SPNs. Therefore, we speculate that an increase in the expression of these transcripts translates into a higher ribosome loading, which is consistent with the observation of increased eIF4E-eIF4G interactions resulting from the loss of FMRP. Alternatively, it may reflect an imbalance in the translation of long versus short length mRNAs^81^.

Our results also show that several transcripts are downregulated in Drd1-SPNs of *Fmr1* KO mice. Among those, *Rgs4* represented an ideal candidate for further investigation. A previous study reported no difference in the expression of *Rgs4* mRNA expression in both the hippocampus and cerebral cortex of *Fmr1* KO mice^82^, further strengthening the striatal-specific role of *Rgs4* in the expression of RRBs. In Drd1-SPNs, endogenous cholinergic signaling through M4Rs promotes LTD of cortico-striatal glutamatergic synapses by suppressing RGS4 activity^48^. At dSPN cortico-striatal synapses, M4R signaling is mediated by RGS4 deactivation, which in turn attenuates mGluR5 signaling through Gαq^83^. M4Rs act as a functional antagonist of cAMP-dependent signaling pathways in Drd1-SPNs^84^ and the strength of dSPN glutamatergic synapses is reciprocally modulated by M4Rs and D1Rs^48^. We hypothesized that reduced expression of *Rgs4* in Drd1-SPNs may underlie the enhanced form of synaptic plasticity occurring at the cortico-striatal synapses of *Fmr1* KO mice (**Fig. 8**). It was therefore surprising that PAM VU0152100 administration corrects the exaggerated cortico-striatal LTD as well as the aberrant RRBs exhibited by *Fmr1* KO mice. Although unexpected, these results highlight the strong association between dysregulated striatal plasticity and RRBs expression, and are consistent with a previous study showing that activation rather than inhibition of M4Rs, which is excessively translated in the hippocampus of *Fmr1* KO mice, corrects core features of FXS, including excessive hippocampal protein synthesis and mGluR-LTD^52^. Therefore, *Rgs4* represents a promising therapeutic target for the modulation of dysregulated translation downstream of mGluR5 in FXS pathological phenotype.

**Figure 8.**
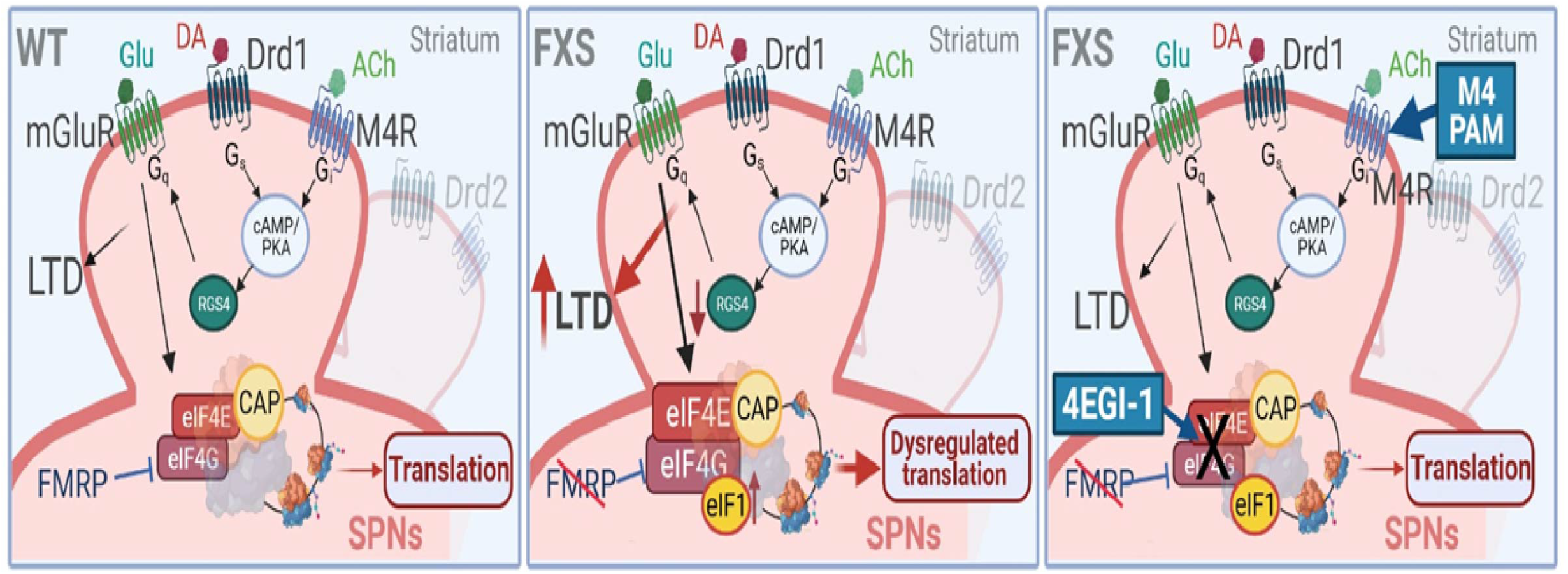
Proposed model for correction of FXS by either 4EGI-1 or M4R PAM VU0152100. Loss of FMRP in FXS leads to enhanced eIF4E-eIF4G interaction resulting in net increase in *de novo* cap-dependent translation in Drd1-SPNs in *Fmr1* KO mice. In FXS, the absence of FMRP leads to the excessive synthesis of eIF1 and downregulated expression of RGS4 therefore cortico-striatal LTD is enhanced in *Fmr1* KO mice. By using 4EGI-1 or M4R PAM VU0152100 LTD and aberrant behavior are rescued. 4EGI-1 also normalizes protein synthesis rate in the striatum of *Fmr1* KO mice.

In sum, our findings support the model (**Fig. 8**) that excessive cap-dependent translation, via increased eIF4E-eIF4G interactions, triggers changes in DLS synaptic composition and function, driving the expression of RRBs displayed by FXS model mice. Our study provides the first evidence of the effect of the disruption of FMRP activity on SPNs synaptic structure and function and the resulting activity imbalance between the direct and indirect pathways. Thus, these circuits may represent a promising therapeutic target for RRBs associated with FXS and ASD, and pharmacologic interventions that remedy striatal dysfunction may assist in the prevention and treatment of this phenotype. Moreover, our study identified new differentially translated mRNAs in dSPNs, providing critical information concerning potential therapeutic targets. Finally, the aberrant synaptic structure/function and we observed should stimulate further investigation on the effect of FMRP loss in the dSPNs and open to the exploration of new therapeutic avenues for the pharmacological modulation of *Rgs4* and other dysregulated transcripts.

## Acknowledgments

We would like to thank Mian Hou and Claudia Farb from Joseph E. LeDoux laboratory for exceptional technical assistance. We thank Dr. Virginia Garcia-Marin for her assistance with deconvolution technique. We are grateful to Maggie Mamcarz for her assistance with mouse breeding and we thank all members of the Klann laboratory for critical feedback and discussions. This study was supported by National Institutes of Health Grants NS034007 (EK), NS047384 (EK), and DP2NS105553 (NXT), U.S. Department of Defense Award W81XWH-15-1-0360 (EK) and Swedish Research Council 2021-02891 (FL).

## Contributions

FL carried out the behavioral experiments, performed slice electrophysiology experiments, and collected and analyzed all *in vivo* and *ex vivo* data. SA carried out and analyzed Trap-Seq. SA, JT, FA and JDZ carried out behavioral experiments and collected *in vivo* and *ex vivo* data. MMO collected *ex vivo* data. PA and CB carried out whole-cell electrophysiology experiments and collect and analyzed whole-cell recordings data. MM and NXT carried out and analyzed the 2-photon Ca2+ imaging data. FL and EK, conceived and designed the studies. SA, AGC, DG, CB and ES participated in the design of the studies. FL and EK designed and coordinated all experiments and wrote the paper. All authors read and commented on the paper.

## Online Methods

### Animals

All procedures involving animals were performed in accordance with protocols approved by the New York University Animal Welfare Committee and followed the National Institutes of Health (NIH) Guide for the Care and Use of Laboratory Animals. All mice were housed in groups of 3–4 animals per cage in the Transgenic Mouse Facility of New York University and maintained in accordance with the US National Institutes of Health Guide for Care and Use of Laboratory Animals^85^.The facility was kept under regular lighting conditions (12 h light/dark cycle) with a regular feeding and cage-cleaning schedule. Room temperature was maintained at 21 ± 2°C. Mice were all maintained on a C57BL/6J genetic background, and all genotypes were determined by polymerase chain reaction (PCR).

#### Fmr1 knockout (Fmr1 KO) mice

*Fmr1* KO mice and their wild-type littermates were bred as previously described^30,86^ and were backcrossed routinely with C57BL/6 mice every three generations.

#### Fmr1 KO/Drd2-EGFP - Fmr1 KO/Drd1a-tdTomato mice

We generated double mutant *Fmr1* KO mice harboring a transgenic BAC containing either the mouse dopamine receptor D1A (*Drd1a*) promoter directing the expression of a modified DsRed fluorescent protein, tdTomato^87^ or the mouse dopamine receptor D2 (*Drd2*) promoter directing the expression of green, fluorescent protein, EGFP^53^. Briefly, *Fmr1* heterozygous female mice were crossed with either *Drd2*-EGFP or *Drd1a*-tdTomato hemizygous BAC transgenic male mice to visualize MSNs of both the direct (striatonigral) and indirect (striatopallidal) pathways (*Drd1a*, direct pathway; Drd2, indirect pathway) in *Fmr1* KO (*Fmr1* KO/*Drd2*-EGFP; *Fmr1* KO/*Drd1a*-tdTomato) and control mice (WT/*Drd2*-EGFP; WT/*Drd1a*-tdTomato).

#### Drd1a-bacTRAP transgenic mice

*Fmr1* KO mice bearing an EGFP-L10a ribosomal fusion protein targeted to the *Drd1* gene were obtaining by crossing *Fmr1* heterozygous female mice with hemizygous bacTRAP transgenic male mice bearing the TRAP transgene (EGFP-L10a) under the control of Drd1a receptor loci in the appropriate BAC (Drd1a-EGFP/Rpl10a; Jackson Laboratory, stock number: 030254)^43^.

#### Fmr1^f/f^ Drd1-Cre transgenic mice

Heterozygous female mice harboring floxed *Fmr1* gene (*Fmr1^f/+^*) generated as previously described^54^ were crossed with heterozygous Drd1-Cre male mouse line obtained from the GENSAT Project at Rockefeller University^53^. Here, a Cre-expression cassette, followed by a polyadenylation sequence to terminate transcription of the fusion transcript immediately after the recombinase gene, was inserted into a BAC vector at the initiating ATG codon in the first coding exon of the gene. The resulting heterozygous male mice (*Fmr1*^f/+^ DAT-Cre) were crossed with *Fmr1*^f/+^ female mice in order to obtain the resulting *Fmr1* conditional knockout (*Fmr1*^f/f^ Drd1-Cre) mice and the respective wild-type (*Fmr1*^+/+^ Drd1-Cre) littermates mice used as a control.

### Stereotaxic surgeries

Surgery for implantation of 26 gauge stainless steel cannulae (Plastics One) was performed as described previously^30^. Briefly, 3-month-old mice mice were anesthetized with a mixture of ketamine (100 mg/kg) and xylazine (10 mg/kg) and mounted on a stereotaxic apparatus. Cannulae were unilaterally implanted in the right lateral ventricle at the following coordinates: −0.22 mm anterioposterior, +1 mm mediolateral, and −2.4 mm dorsoventral (Santini et al., 2017). Mice were given at least 1 week after surgery to recover.

### Drug preparation

4EGI-1 was prepared as described previously^30,34,88^. 4EGI-1 (#324517, Merck-Millipore) was dissolved in 100% DMSO and diluted in vehicle (0.5% (2-hydroxypropyl)-b-cyclodextrin and 1% DMSO in artificial CSF) to a final concentration of 100 μM. 4EGI-1 (1 μl; 20 μM) was infused intracerebroventricularly at a rate of 0.5 μl/min; injectors remained in the guide cannula for 3 min after the infusion. The M4 muscarinic receptor (M4R) positive allosteric modulator (PAM) VU0152100 (#V5015, Sigma-Aldrich) was dissolved in 10% DMSO + 10% Tween-80 in PBS and injected intraperitoneally (i.p.) at the dose of 56 mg/kg^52^. Control mice received equivalent volume of vehicle solutions. Both drugs and vehicles were administrated 1h prior behavioral experiments.

### Behavior

Mice were acclimated to the testing room 30 min prior to each behavioral experiment and all behavioral apparatuses were cleaned between each trial with 30% ethanol. All behavior sessions were conducted during the light cycle and mice were randomly assigned for experimental conditions including drug or vehicle infusions, and for the order of testing in any given experimental paradigm. Experimenter were blind to genotype and experimental conditions while performing and scoring all behavioral tasks.

### Pole test

The pole test (modified from^89^) was used to assess striatal-based motor dysfunction in mice. Mice were placed at the top of a 50 cm vertical pole with a diameter of 1 cm and a triangular base stand. The pole was placed in the home cage to favour mice descent from the pole. Recording started when the animal began the turning movement to descend. The total time that mice spent to descend into the cage (T_total_; sec) and to turn themselves downward (Latency to turn, T_turn_; sec) were recorded. Mice were subjected to a 3-trial training session where they were trained to turn around and descend the pole followed, 30 minutes later, by the testing session. The test was video-recorded, and the performance was scored manually. A maximum score of 20 sec (cut-off) was assigned to a mouse that fell off from the pole.

### Open field test

The open field (OF) test was used to measure the spontaneous general locomotor activity and anxiety-like behavior and was performed as previously described^34,90^. The total distance travelled was recorder over a 15 min period by using a computerized video tracking system (Activity Monitor software for OF). The data were pooled according to genotype, and a mean value was determined for each group.

### Novel home cage test

The novel home cage (NHC) test was used to assess the spontaneous horizontal motor activity as novelty-induced exploratory response^90,91^. Mice were placed in a 35 × 22 × 22 cm experimental cage with the floor covered with bedding. Locomotor activity (expressed in cm) was recorded over a 60 min period by using a computerized video tracking system (Noldus, EthoVision XT). The parameter tested was the total distance travelled during the test.

### Cylinder test

The cylinder test (modified from^92^) was used to assess the vertical motor activity. Briefly, mice were placed in an open-top, clear glass cylinder (diameter: 13.6 cm, height: 17.2 cm), and allowed to explore by rearing and touching the walls of the cylinder with their forelimb paws. Motor performance was recorded for 3 min and the time spent rearing (sec), and the number of rearing were analysed.

### Drag test

The drag test was performed as previously described^90,93^ and gives information regarding the time to initiate (akinesia) and execute (bradykinesia) a movement. Briefly, mice were lifted from the tail (allowing the forepaws to rest on the table) and dragged backwards at a constant speed (~20 cm/s) for a fixed distance (100 cm). The number of steps made by each forepaw was recorded. Five determinations were collected for each animal. The test was performed on two consecutive days.

### Self-grooming behavior

To test repetitive self-grooming behavior, mice were individually placed in clean empty cages without bedding for a period of 60 min under conditions of white noise. During the first 50 min mice were allowed to habituate to the empty cage. Cumulative time spent in spontaneous repetitive grooming behavior was scored during the last 10 min (modified from^94^).

### Marble burying test

The marble burying (MB) test was performed as previously described^34^. Briefly, mice were placed individually in clean cages containing fresh bedding (5 cm deep) and 20 black marbles arranged in five evenly spaced rows of four marbles each. Testing consisted of a 30 min period under white noise conditions. The number of marbles buried at the end of this period was recorded as measure for repetitive behavior.

### Nestlet shredding test

The nestlet shredding (NS) test was performed as previously described^95^ and used to assess repetitive behavior. Briefly, mice were place individually in clean cages containing fresh bedding (0.5□cm deep), and one commercially available preweighed cotton fiber (nestlet) (5□cm□×□5□cm, 5□mm thick, ~2.5□g) in each test cage. Mice were left undisturbed in the cage with the nestlet for 30 min. After test completion remaining intact nestlet material was removed from the cage with forceps and allow to dry overnight. The remaining unshredded nestlet was weighed, and the weight difference was divided by the starting weight to calculate percentage of nestlet shredded. Food and water were withheld during the test.

### Immunoprecipitation

Pull-down assay was performed in 3 to 4-month-old mice as previously described^30^. Striata were dissected 1 hour after intracerebroventricular infusions with 4EGI-1 (100 μM) or an equivalent volume of vehicle and, flash frozen on dry ice. Tissues were sonicated in cold lysis buffer containing 150 mM NaCl, 10 mM MgCl_2_, 30 mM tris buffer (pH 8.0), 1 mM DTT, 1.5% Triton X-100, protease and ribonuclease inhibitors (10 μl/ml). 500 μg of lysate were incubated with 30 μl of m^7^GTP beads (#AC155; Jena Bioscience) for 1 hours at 4°C. The beads were centrifuged for 1min at 6000 rpm, and the supernatant was collected. The beads were then washed three times in wash buffer containing 100 mM KCl, 50 mM tris buffer (pH 7.4), 5 mM MgCl_2_, 0.5%Triton X-100]. Finally, the beads were eluted with 5X Laemmli buffer and analyzed on western blotting. The following antibodies were used in the western blotting analysis: rabbit anti-eIF4E (#A301-153A; Bethyl Laboratories; 1:1000), rabbit anti-eIF4G (#C45A4; Cell signalling; 1:1000) and rabbit anti-FMRP (#834601; Biolegend, 1:500).

### Surface labelling of *de novo* protein synthesis (SUnSET)

A protocol adapted from the SUnSET method was used to label newly synthetised proteins as previously described^34,90,96^. Briefly, 400 μm-thick coronal striatal slices of the brain of 3- to 4- month-old *Fmr1* KO and control mice were prepared using a vibratome. Slices were allowed to recover in aCSF at 32 °C for 1 h and subsequently treated with puromycin (#P8833, Sigma-Aldrich, 5 μg/mL) for 45 mins. For slices subjected to pharmacological pretreatment, anisomycin (#1290, Tocris, 20 μM) and 4EGI-1 (100 μM) were added to aCSF 30 min prior to puromycin treatment. Newly synthesized proteins were end-labeled with puromycin. Striatum was micro-dissected from the brain slices and flash frozen on dry ice and lysed. 40 μg of puromycylated protein lysates were analysed on western blotting. Protein synthesis levels were determined by taking total lane density in the molecular weight range of 10–250 kDa. Comparisons of protein synthesis levels between both genotypes were made by normalizing to the average WT signal.

### Western blotting

Dorsolateral striatum was micro-dissected from the brain slices of 3 to 4-month-old *Fmr1* KO and WT mice and sonicated in ice-cold homogenization buffer (10mM HEPES, 150mM NaCl, 50 mM NaF, 1 mM EDTA, 1 mM EGTA, 10 mM Na_4_P_2_O_7_, 1% Triton X-100, 0.1% SDS and 10% glycerol) that was freshly supplemented with HALT protease and phosphatase inhibitor cocktail (#78441; Thermo Scientific; 1/10 total volume). Aliquots (2 μl) of the homogenate were used for protein determination with a BCA (bicinchoninic acid) assay kit (GE Healthcare). Samples were prepared with 5X sample buffer (0.25M Tris-HCl pH6.8, 10% SDS, 0.05% bromophenol blue, 50% glycerol and 25% − β mercaptoethanol) and heat denatured at 95 °C for 5 min. 40 μg protein per lane was run in pre-cast 4–12% Bis-Tris gels (Invitrogen) and subjected to SDSPAGE followed by wet gel transfer to polyvinylidene difluoride (PVDF; Immobilon-Psq, Millipore Corporation, Billerica, USA) membranes. Membranes were blocked for 90 min with 5% milk in Tris-buffered saline supplemented with 0.1% Tween-20 (TBST) and then were probed overnight at 4 °C using mouse anti-puromycin primary antibodies (#MABE343; Millipore; 1:1000). Membranes were probed with horseradish peroxidase-conjugated secondary IgG (Promega; 1:7000) for 1 h at room temperature. Signals from membranes were detected with ECL chemiluminescence (GE Healthcare Amersham™) using Alpha Imager 3.4 software and the FluorChem Protein Simple instrument. Membranes were then stripped, reblocked probed with rabbit anti-FMRP (#834601; Biolegend; 1:500) and rabbit anti-GAPDH (#2118; Cell Signalling; 1:1000) primary antibody. The anti-GAPDH antibody was used to estimate the total amount of protein. Membranes were imaged for the respective antibodies again as described. Exposures were set to obtain signals at the linear range and then normalized by total protein and quantified via densitometry using ImageJ software (NIH, USA).

### Fluorescent labelling of *de novo* protein synthesis (FUNCAT)

FUNCAT method was used to label *de novo* protein synthesis in Drd1- or Drd2-MSNs and it was performed as previously described^90^ with minor modifications. Briefly, 400 μm coronal striatal slices from *Fmr1* KO/Drd2-EGFP or Drd1a-tdTomato BAC transgenic male mice and their littermate controls were incubated with azidohomoalanine (AHA) at 32 °C for 2.5 h. For slices subjected to pharmacological pretreatment, 4EGI-1 (100 μM) were added to aCSF 30 min prior to AHA incubation. At the end of the incubation slices were fixed overnight at 4 °C in 4% PFA and, re-sliced using a vibratome (Leica VT1200S; Leica Microsystems; Bannockburn, IL) to a thickness of 30 μm. Free floating sections were collected in Tris-buffered saline (TBS), blocked and permeabilized with 5% bovine serum albumin, 5% normal goat serum (NGS), 0.3% Triton-X-100 in TBS for 90 min (at RT). Overnight cycloaddition was performed on slices by using cyclo-addition reaction mix (Click-iT™ Cell Reaction Buffer Kit, Invitrogen, Ltd, Paisley, UK) at 4 °C with gentle rocking. For slices expressing Drd1a-tdTomato MSNs AHA was detected using an Alexa Fluor™ 488 Alkyne, Triethylammonium Salt (Invitrogen, Carlasbad, CA, USA), whereas for slices expressing Drd2-EGFP an Alexa Fluor™ 647 Alkyne, Triethylammonium Salt (Invitrogen, Carlasbad, CA, USA) was used. Slices were then probed with primary antibodies: rabbit anti-RFP (#600-401-379; Rockland; 1:500), or chicken anti-GFP antibody (#ab13970; abcam; 1:500) for 3 hours at room temperature. Slices were then rinsed in TBS and incubated with either Alexa Fluor™568 goat anti-chicken (1:400) or Alexa Fluor™488 goat anti-chicken (1:400) secondary antibody (Invitrogen, Carlasbad, CA, USA). Finally, slices were rinsed with TBS and mounted using DAPI fluoromount-G™ (Electron Microscopy Sciences, Hatfield, PA, USA) and processed for fluorescence imaging using Leica LSM8 confocal microscope. Images were obtained using the same settings for all samples within an experiment. Fluorescence was quantified using ImageJ software (NIH, USA) as previously described^90,97^.

### Dendritic spine density

Dendritic spine density analysis was performed as previously described^34,42^. To visualize dendrites and dendritic spines we collected coronal cortico-striatal slices (200 μm) from double mutant Fmr1 KO mice harboring a transgenic BAC containing either the mouse dopamine receptor D1A (Drd1 a) promoter directing the expression of a modified DsRed fluorescent protein, tdTomato^87^ or the mouse dopamine receptor D2 (Drd2) promoter directing the expression of green, fluorescent protein, EGFP^53^. Images were acquired by generating maximum intensity projections from z-stacks using Leica LSM8 confocal microscope. Images were then subjected to deconvolution technique using a blind deconvolution package from Huygens Professional (Scientific Volume Imaging, The Netherlands). To quantify, we identified a 20-30 μm dendritic segments that were ≥ 5 μm distant from the proximal and the distal and counted individual spines using ImageJ.

### Translating ribosome affinity purification (TRAP)

Trap-Seq was carried out as described in^98^. Briefly, striata from WT (n=6) and *Fmr1* KO (n=5) mice expressing the EGFP-tagged Rpl10a protein in dopamine receptor Drd1-expressing medium spiny neurons were lysed by dounce homogenization (40 strokes) in 25 mM Hepes-HCl (pH 7.3), 150 mM KCl, 10 mM MgCl_2_, 0.5 mM DTT, 100 μgml^−1^ Cycloheximide, 10 μlml^−1^ RNasin (Promega, Madison, WI) and 10 μlml^−1^ Superase-In (Life Technologies) RNase inhibitors, and 1X Halt protease/phosphatase inhibitor on ice. Nuclei were pelleted by centrifuging the lysates at 2000g. NP-40 (Sigma) and DHPC (Avanti Polar Lipids, Alabaster, AL) were each added to the supernatant to a final concentration of 1%, following which the lysates were centrifuged at 20,000g in order to pellet insoluble membranes. A small aliquot of the supernatant was saved for RNA-Seq (input). EGFP-tagged ribosomal protein L10a was precipitated by incubating the remaining supernatant overnight (4°C) with 100 μg of monoclonal anti-EGFP antibodies (50 μg each of clones 19C8 and 19F7) bound to biotinylated-Protein L (Pierce, Thermo Fisher, Waltham, MA) coated streptavidin-conjugated magnetic beads (Life Technologies). The magnetic beads were then washed four times in high-salt buffer consisting of 10 mM Hepes-HCl (pH 7.3), 350 mM KCl, 5 mM MgCl_2_, 1% NP-40, 0.5 mM DTT, 100 μgml^−1^ Cycloheximide, and RNasin and Superase-In RNase inhibitors (Promega). Bound RNA was eluted and purified using the Absolutely RNA Nanoprep kit (Agilent, Santa Clara, CA). Sequencing libraries (non strand-specific) from the IP and input RNA were prepared using the Nugen Ovation Trio Low Input RNA kit at the NYUMC Genome Technology Center. Libraries were sequenced on the Illumina NovaSeq S1 100 Cycle Flow Cell to generate 50-cycle paired-end reads. All pulldowns and sequencing were carried out in a single batch. Reads were aligned to mm10 with the *STAR* aligner^99^. Reads mapped to genes annotated in the Gencode primary assembly were counted during alignment. Differential expression analyses were carried out using *DESeq2*^100^. The R package *fgsea* was used to carry out gene set enrichment analyses. The *phyper* function in R was used to carry out hypergeometric tests to calculate the probability that genes significantly altered in ribosome association in D1 neurons overlapped by chance with those altered in striatal mRNA expression. Canonical mRNA CDS lengths were obtained from Ensembl, log2 transformed, and its histogram divided into regular intervals in order to bin mRNAs into 6 CDS length bins. To divide mRNAs into 50 bins evenly, the total number of robustly expressed mRNAs (counts per million > 1 for all samples) were divided by 50 and the number of genes indicated by the remainder was removed at random. For example, 33 genes were removed at random from 12033 genes to ensure that each length bin contained 240 mRNAs.

### Slice preparation

Coronal striatal sections (300 μm) from mice 3-4 months of age were prepared as described previously^34,90^. Ice cold oxygenated (95% O_2_/5% CO_2_) cutting solution containing the following (in mM): 110 Sucrose, 60 NaCl, 3 KCl, 1.25 NaH_2_PO_4_, 28 NaHCO_3_, 0.5 CaCl_2_, 7 MgCl_2_, 5 Glucose was used for slice isolation. Then, slices were transferred to oxygenated artificial cerebrospinal fluid (ACSF) containing the following (in mM): 125 NaCl, 2.5 KCl, 1.25 NaH_2_PO_4_, 25 NaHCO_3_, 25 D-glucose, 2 CaCl_2_, and 1 MgCl_2_. Slices were incubated at room temperature and then were placed in the recording chamber for additional recovery time of 60 min at 32°C. For bath application the drugs were made and stored as concentrated stock solutions and diluted 1000-fold when applied to the perfusate. For whole-cell recordings, mice (3-4 months old) were anesthetized with isoflurane and intracardially perfused with ice cold cutting solution containing the following (in mM): 65 sucrose, 76 NaCl, 25 NaHCO_3_, 1.4 NaH_2_PO_4_, 25 glucose, 2.5 KCl, 7 MgCl_2_, 0.4 Na ascorbate, and 2 Na pyruvate (bubbled with 95% O_2_/5% CO_2_). 300 μm coronal sections were cut in cutting solution before being transferred to ACSF containing the following (in mM): 120 NaCl, 25 NaHCO_3_, 1.4 NaH_2_PO_4_, 21 glucose, 2.5 KCl, 2 CaCl_2_, 1 MgCl_2_, 0.4 Na ascorbate, and 2 Na pyruvate (bubbled with 95% O_2_/5% CO_2_). Slices were recovered for 30 min at 35°C and subsequently stored at 24°C for at least 30 min. All slice recordings were conducted at 30-32°C.

### Electrophysiology

Extracellular field excitatory postsynaptic potentials (fEPSPs) were recorded as described previously^34,90^. Briefly, coronal striatal slices from mice were isolated and transferred to recording chambers (preheated to 32 °C), where they were superfused with oxygenated ACSF. In all the experiments, baseline synaptic transmission was monitored for at least 20 min before long-term depression (LTD) induction. Three trains of high-frequency stimulation (HFS; 3 s duration, 100 Hz frequency at 20 s intervals) were used to induced LTD in striatal slices. After induction of striatal LTD, fEPSPs were collected for an additional 70 min. Slices were treated with either 4EGI-1 (100 μM), VU0152100 (5 μM) or vehicle applied 10 min before the tetanus and perfused for 70□min after tetanus. Both drugs and vehicles were maintained in the bath for the duration of the recordings. Slope values of fEPSP were expressed as a percent of the baseline average before LTD induction and were acquired using pClamp 10 (Axon Instruments, Foster City, CA). Data collection and analysis were not performed blind to the conditions of the experiments.

Targeted whole-cell recordings were made from SPNs in the dorsolateral striatum using infrared-differential interference contrast. Drd1+ and Drd2+ SPNs were identified using fluorescent illumination in tomato +/- cells in Drd1 tdtomato mice or GFP +/- cells in Drd2 EGFP mice, as described previously^35,36^. Recordings were made from mutant male mice and age-matched male littermates. Voltage clamp experiments were made with borosilicate pipettes (3 5 MΩ) filled with the following (in mM): 135 Cs gluconate, 10 HEPES, 10 Na phosphocreatine, 4 Mg2 ATP, 0.4 NaGTP, 10 TEA, 2 QX-314, and 10 EGTA, pH 7.3 with CsOH (290 295 mOsm). In all experiments, 10 μM Gabazine was used to block GABA receptors and 1 μm TTX was included to block action potentials. Physiology data were acquired using National Instruments boards and custom software written in MATLAB (MathWorks). mEPSC measurements and quantification were performed using the NeuroMatic plugin for Igor Pro. The minimal threshold for detection was 2 pA and mEPSCs were analyzed across a minimum of 20 seconds of recording.

### Data acquisition and analysis

All data are presented as the mean ± SEM. Data were analyzed using GraphPad Prism 8. For behavioral experiments the experimenter was blinded to the genotype of the animals during behavioral testing. For two-group comparisons, statistical significance was determined by parametric and nonparametric two-tailed Student’s t tests or Mann-Whitney test. Multi-groups were analyzed using one-way ANOVA or two-way ANOVA. P values < 0.05 were considered statistically significant. Extreme outliers were detected by applying Grubbs’ method with α = 0.05 to each experimental group and eliminated from further analysis (GraphPad software). Sample size was chosen following previous publications. Data distribution was assessed to be normal. Variance was similar between the groups that were being statistically compared based on our observation.

### *In vivo* striatal imaging

Mice were implanted with a custom titanium headpost (Parkell) and stereotaxically (Kopf Instruments) injected with AAV1-Syn-GCaMP6f-WPRE-SV40 virus (University of Pennsylvania Vector Core) into the right dorsolateral striatum at four different locations (from bregma, AP/ML = 0.7/1.7 mm, 0.7/2.3 mm, 1.3/1.7 mm and 1.3/2.3 mm) while under anesthesia (1.5% isoflurane in oxygen). 100-200 nl virus was injected at each location (DV = −1.8 mm from dura) at a rate of 100 nl.min^−1^ using pulled glass capillaries (Drummond) connected to a 5 μl Hamilton syringe pump (KD Scientific). A 3 mm craniotomy was subsequently performed, as previously described^38^. Cortical tissue was removed with suction until the external capsule above dorsal striatum surface was exposed and a custom cannula was lowered above striatum and permanently cemented to the skull using C&B metabond^38^. Two weeks after surgery, mice were gradually introduced to the recording setup and trained to spontaneously locomote while head-fixed on a circular treadmill (Ware Manufacturing) mounted on a rotary encoder (MA3-A10-125-B; US Digital) under a resonant galvanometer two-photon microscope (Bergamo II, Thorlabs). Imaging sessions began as soon as mice reliably and comfortably engaged in spontaneous bouts of locomotion for at least 30 min. Striatal fields of view (~500 μm x 500 μm per field) with fluorescence for both GCaMP6f and tdTomato were acquired at 30 Hz (ScanImage, Vidrio Technologies) using 940 nm excitation light (Chameleon Vision II, Coherent; ~100 mW at sample) through a 20X air objective (#58373, Edmund Optics).

Acquired movies were processed using MATLAB scripts generously provided by the Harvey Lab (https://github.com/HarveyLab/Acquisition2P_class.git) to correct for movement artefacts, semi-automatically segment SPNs and extract calcium fluorescence after neuropil subtraction. dSPNs were distinguished from iSPNs manually based on tdTomato fluorescence. Individual calcium transients were detected using MATLAB’s ‘findpeaks’ function on each cell’s ΔF/F trace smoothed with a 170 ms sliding window. Individual neurons were deemed ‘active’ if they displayed at minimum one calcium transient per imaging session. Imaging and behavioral data were quantified in MATLAB by using custom-code available online (https://github.com/TritschLab/TLab-2P-analysis)^38^.

## Data and Software Availability

Raw RNA-Seq sequencing reads for the immunoprecipitated Drd1-SPN EGFP-L10a copurifying RNA (IP) and the striatal lysate RNA (total) from WT and FXS mice striata are publicly available at NCBI GEO under the accession number GSE165872. The data that support the findings of this study are available from the corresponding author upon request.

---

**Supplementary Figure 1.**
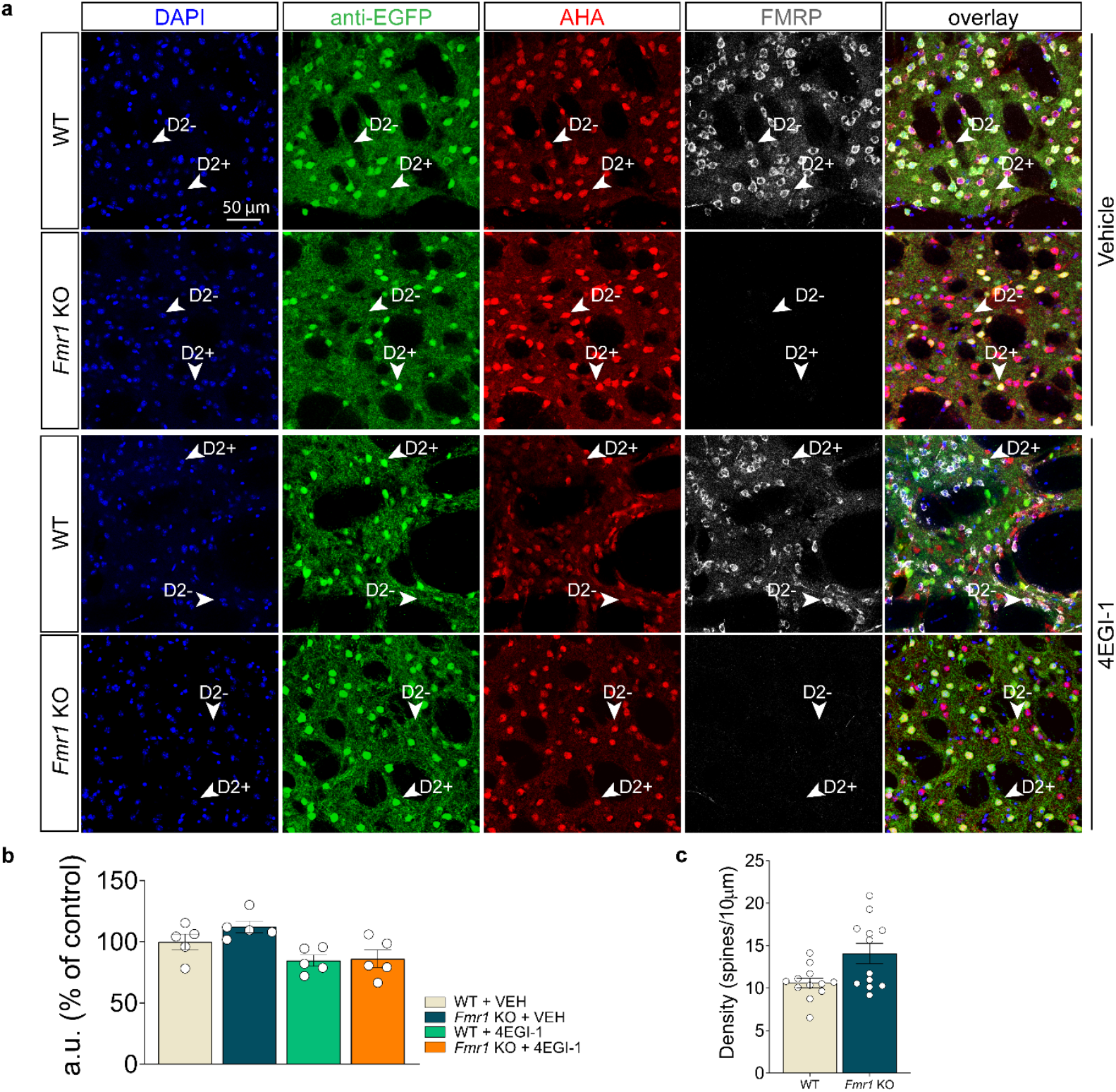
Lack of *Fmr1* does not results in dysregulated *de novo* translation in Drd2-SPNs. **a**, Representative DLS immunofluorescence images of DAPI (blue), anti-EGFP (green), anti-FMRP (grey) and incorporation of AHA (red) detected by FUNCAT with alkyne-Alexa 647 in cortico-striatal slices from *Fmr1* KO/Drd2 EGFP BAC transgenic mice and their WT littermates (scale bar represents 50 μm) treated with VEH (first two rows from the top) or 4EGI-1(last two rows from the top). **b**, Quantification of increased AHA-alkyne-Alexa 647 signal in fluorescent arbitrary units (a.u.) expressed as % of control in Drd2-SPNs (anti-EGFP+ neurons; green) from DLS *Fmr1* KO/Drd2 EGFP BAC transgenic mice and their WT littermates. Cell soma intensity was measured in ImageJ software (FIJI). Statistical significance was determined by using two-way ANOVA followed by Bonferroni’s multiple comparisons test (genotype x treatment interaction: *F_(1,16)_*= 0.85, *P*=0.37; genotype: *F_(1,16)_*= 1.42, *P*=0.25; treatment: *F_(1,16)_*= 12.79, \*\**P*<0.01). Data are shown as mean ± s.e.m. of *n* = 5/6 mice per group (average of *n* = 20 somas per slice, *n* = 2 slices per mouse, from three independent experiments). **c**, *Fmr1* KO mice show no significant difference in overall dendritic spine density in DLS (Mann-Whitney test, *P*=0,052).

**Supplementary Figure 2.**
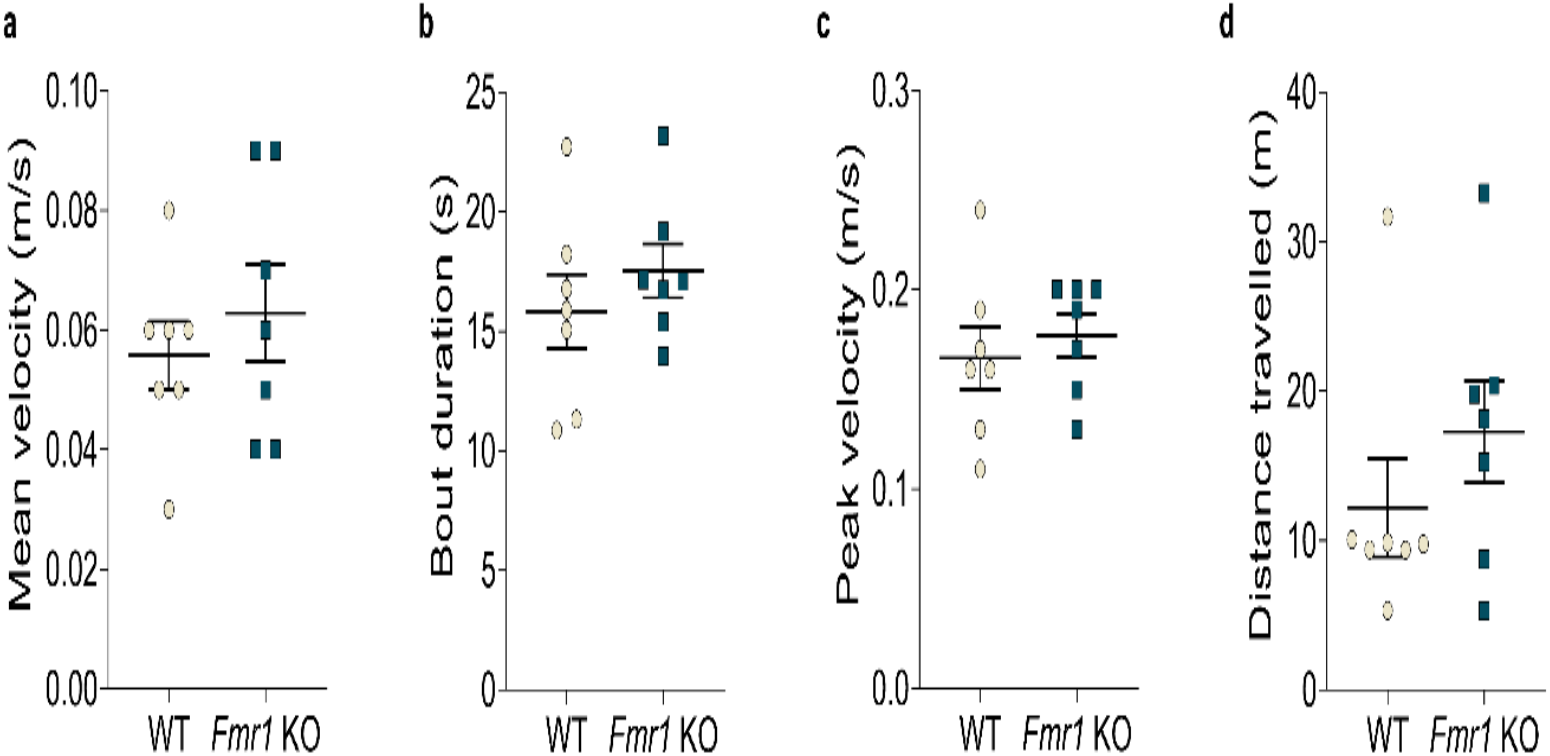
Self-initiated forward locomotion on treadmill during *in vivo* two-photon imaging. **a**, Summary plot of mean self-initiated treadmill velocity in WT (*n*= 4) or *Fmr1* KO (*n*= 4) mice (*P* = 0.63; Mann-Whitney test, *n*=7 imaging sessions per genotype). **b**, Same as (a) for locomotor bout duration (*P*= 0.38; Mann-Whitney test, *n*=7 imaging sessions per genotype). **c**, Same as (a) for locomotor peak velocity (*P* = 0.40; Mann-Whitney test, *n*=7 imaging sessions per genotype). **d**, Same as (a) for total distance travelled (*P* = 0.38; Mann-Whitney test, *n*=7 imaging sessions per genotype).

**Supplementary Figure 3.**
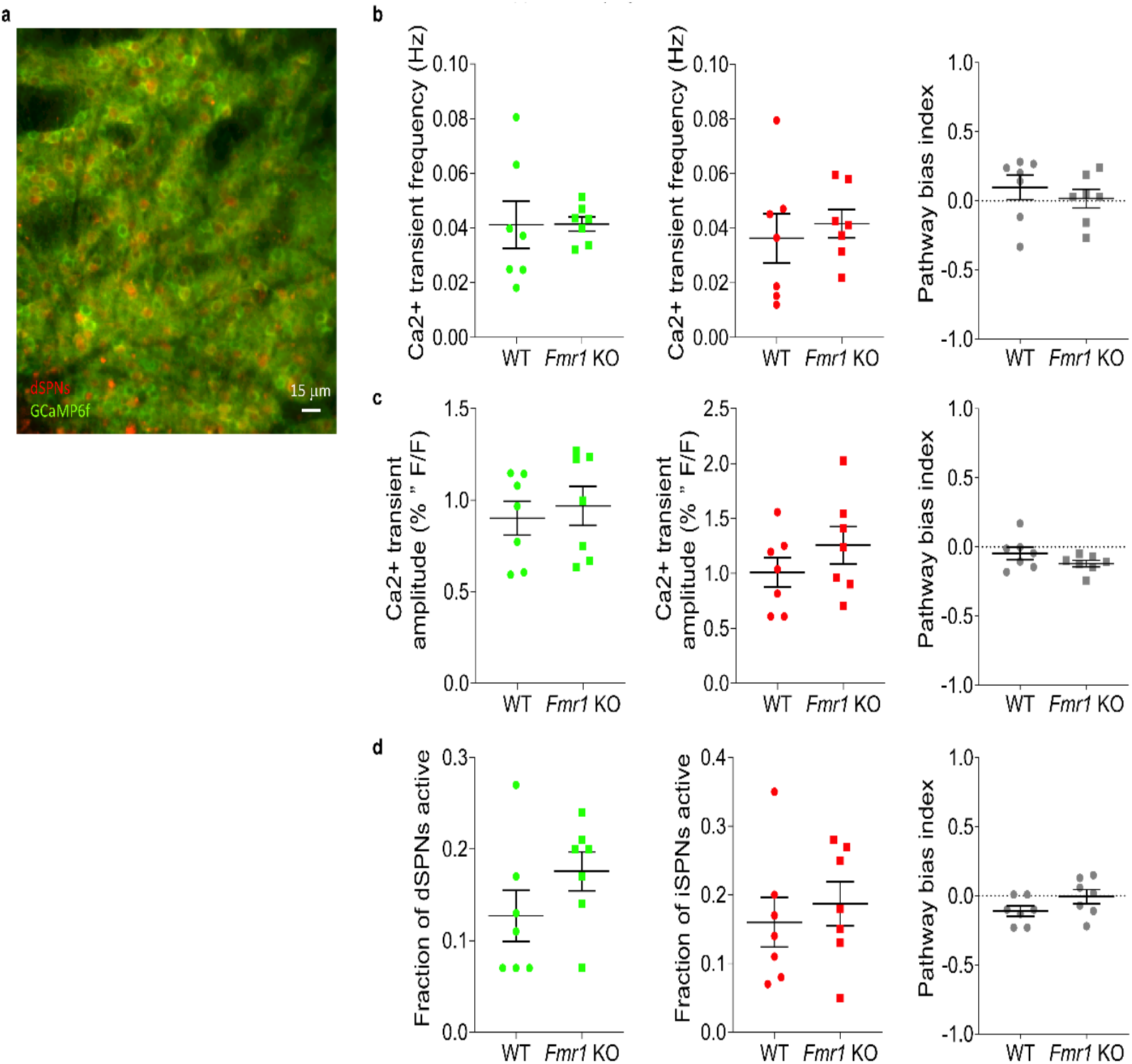
*In vivo* two-photon Ca2+ imaging of dSPNs in the DLS of WT and *Fmr1* KO mice. **a**, Representative two-photon maximum projection image of dorsal striatum. Red: tdTomato-labeled dSPNs; green: striatal neurons expressing GCaMP6f (scale bar: 15 μm). **b**, Mean Ca2+ transient frequency per dSPN (left, green) and iSPN (middle, red) active during self-initiated movement in each of 7 FOVs from 4 mice per genotype. Right: mean Ca2+ transient frequency bias index. **c**, Same as (b) for mean amplitude of Ca2+ transients in active dSPNs and iSPNs. **d**, Same as (b) for the total fraction of all imaged dSPNs or iSPNs that exhibit Ca2+ transients.

**Supplementary Figure 4.**
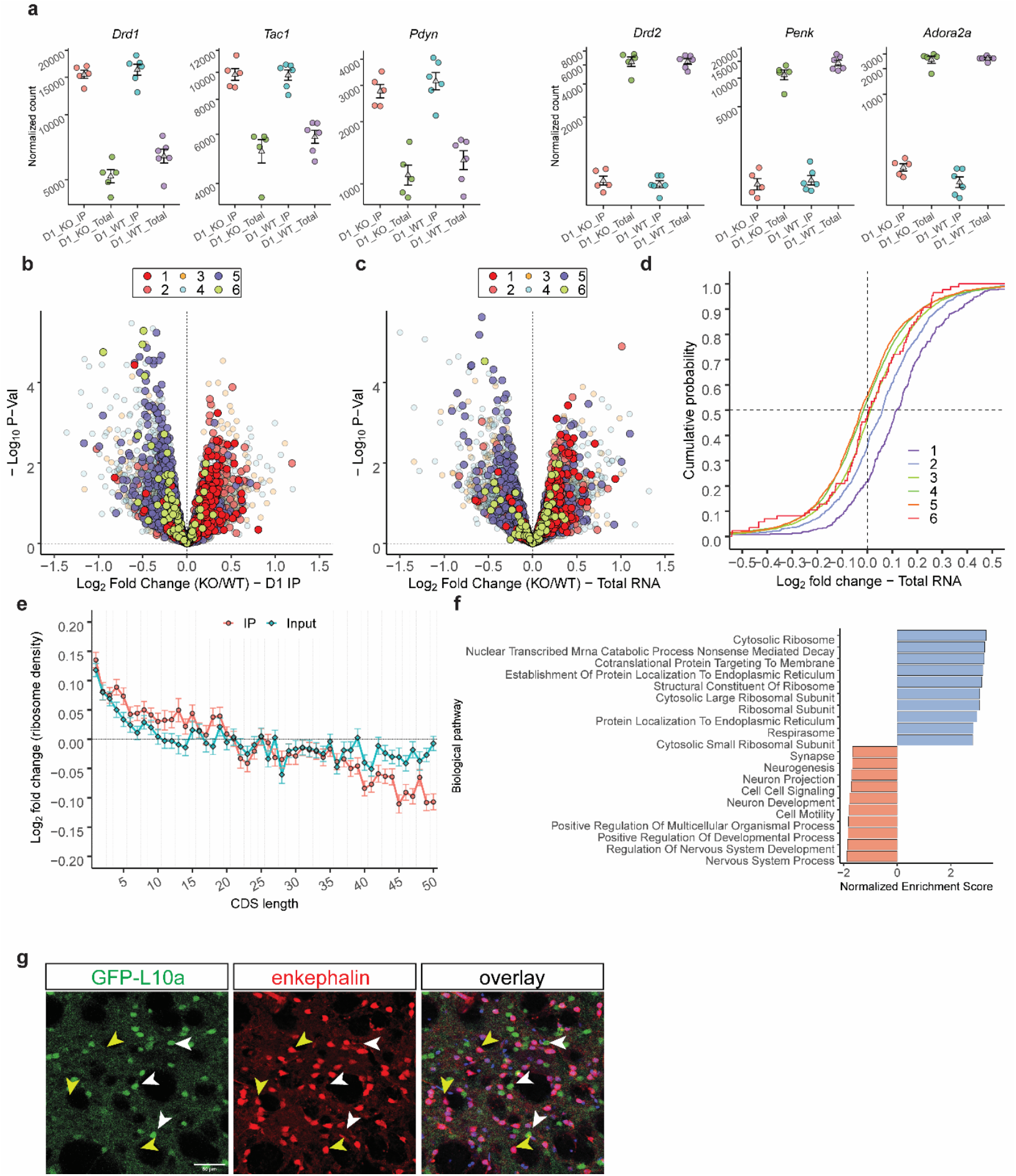
Dysregulated mRNA in Drd1-SPNs of *Fmr1* KO mice. **a**, Trap-Seq of Drd1-SPNs reveals an enrichment of D1 markers and re-duction of markers of Drd2-SPNs. Characteristic markers of Drd1-SPNs including dopamine receptor D1 (Drd1), substance P (Tac1), and dynorphin (Pdyn) are enriched in the IP, while dopamine receptor D2 (Drd2), adenosine 2a receptor (Adora2a), and enkephalin (Penk), which are characteristic markers of D2 MSNs, are reduced relative to their overall mRNA expression in the striatum, in both WT and FXS mice. **b**, Significance (unadjusted (nominal) p-value) vs. log2-fold-change in ribosome association (IP) in Drd1-SPNs and **c**, overall striatal mRNA expression (Total) between FXS and WT mice. Messenger RNAs are divided into six bins in ascending order of their CDS lengths, with bin 1 harboring mRNAs with the shortest CDSs, and bin 6 the longest. mRNAs are color-coded by their CDS length bins. mRNAs with long CDSs are enriched in genes showing significant reduction in ribosome association in D1 MSNs of mice lacking FMRP, while those with the shortest CDSs are over-represented in genes exhibiting significant increase in ribosome association. A similar but weaker trend is also observed in alterations in overall RNA expression in the striata of FXS model mice. **d**, Cumulative distribution of log2-fold-changes (FXS/WT) in RNA expression in the striatum of FXS model mice, as a function of CDS length. mRNAs with the shortest CDSs are especially likely to show elevated expression in the striatum. **e**, Comparison of log2-fold-changes in ribosome association in Drd1-SPNs of FXS, and overall RNA expression in the striatum, by CDS length. mRNAs are divided into 50 bins, with each containing 201 mRNAs. mRNAs with the shortest CDSs exhibit increased ribosome association in D1 neurons of FXS model mice, while those with the longest CDSs exhibit reduced ribosome association. A positive-to-negative gradation is observed with CDS length in log2-fold-changes in ribosome association. LFCs in overall mRNA expression of short CDS mRNAs in the striata of mice lacking FMRP closely track the LFCs in ribosome association in D1 MSNs of FXS model mice. For long CDS mRNAs, much larger and overwhelmingly negative alterations in ribosome association in Drd1-SPNs are observed compared to overall mRNA expression in the striata. **f**, Enrichment scores of the top 10 gene ontologies (GOs) enriched in the WT or FXS striata, determined by GSEA on genes ranked by their fold changes RNA expression. mRNAs coding for ribosomal proteins are overabundant in FXS striata, while GOs associated to synapse and glutamatergic signaling are reduced. **g**, Confocal images show selective expression of GFP-L10a in Drd1-SPNs verified at the protein expression level by immunostaining coronal slices containing the DLS with enkephalin antibody (red), a marker for Drd2-SPNs. EGFP-L10a expression did not costained with enkephalin in DLS Drd1-SPNs of Drd1a-bacTRAP transgenic mice (scale bar represents 50 μm). White arrows indicate Drd1-SPNs (green) and enkephalin (red) staining; yellow arrows indicate non-Drd1-SPNs and enkephalin (red) co-staining.

